# System-level characterization of engineered and evolved formatotrophic *E. coli* strains

**DOI:** 10.1101/2024.11.11.622960

**Authors:** Suzan Yilmaz, Boas Kanis, Rensco A. H. Hogers, Sara Benito-Vaquerizo, Jörg Kahnt, Timo Glatter, Beau Dronsella, Tobias J. Erb, Maria Suarez-Diez, Nico J. Claassens

## Abstract

One-carbon compounds, such as formate, are promising and sustainable feedstocks for microbial bioproduction of fuels and chemicals. Growth of *Escherichia coli* on formate was recently achieved by introducing the reductive glycine pathway (rGlyP) into its genome, which is theoretically the most energy-efficient aerobic formate assimilation pathway. While adaptive laboratory evolution was used to enhance the growth rate and biomass yield significantly, still the best performing formatotrophic *E. coli* strain did not approach the theoretical optimal biomass yield of the rGlyP. In this study, we investigated these previously engineered formatotrophic *E. coli* strains to find out why the biomass yield was sub-optimal and how it may be improved. Through a combination of metabolic modelling, genomic and proteomic analysis, we identified several potential metabolic bottlenecks and future targets for optimization. This study also reveals further insights in the evolutionary mutations and related changes in proteome allocation that supported the already substantially improved growth of formatotrophic *E. coli* strain. This systems-level analysis provides key insights to realize high-yield, fast growing formatotrophic strains for future bioproduction.

## 1. Introduction

Emissions of CO_2_ originating from fossil resources are a global problem leading to a climate crisis with potentially disastrous consequences [1], [2]. A promising alternative to fossil-based production methods is microbial bioproduction of value-added chemicals and fuels. However, current microbial bioproduction processes often rely on agriculture-intensive feedstocks like glucose, which compete for arable land and resources with food production, and thus cannot be scaled to the extend required to replace fossil resources. Therefore, one-carbon compounds such as formate, methanol, methane, and carbon monoxide are being explored as sustainable feedstocks for microbial bioproduction [3], [4], [5]. Formate and methanol are particularly of interest due to their high solubility in water [6], [7]. These compounds can be produced directly (formate) or in two steps (methanol) by electrochemical reduction of CO_2_ using electricity derived from renewable resources. In this work we focus on formate as a feedstock.

Some natural formate-utilizing microbes have been identified, but few have been well-characterized, and they generally lack effective and robust genetic engineering tools [7]. Consequently, optimization of metabolic pathways in most natural formatotrophs is difficult and engineering of novel (production) pathways is often not feasible. In order to explore beyond the limited product spectrum offered by natural formatotrophs, formate assimilation pathways can instead be introduced in model hosts like *Escherichia coli*. Several natural and synthetic formate assimilation pathways have been described, such as the Calvin Benson Bassham (CBB) cycle, the Serine Cycle, the Wood-Ljungdahl Pathway, the Serine-Threonine Cycle, and the reductive Glycine Pathway (rGlyP) [8], [9]. The rGlyP is the most energy efficient aerobic formate assimilation pathway [6] and was first designed by Bar-Even et al. in 2013 [10] and later discovered in the natural formatotroph and autotroph *Desulfovibrio desulfuricans* [11]. Soon after design of the pathway, efforts to engineer the rGlyP in *E. coli* started through a step-wise, modular engineering approach [12], [13], [14], [15].

The rGlyP in *E. coli* consists of an energy generation reaction and a formate assimilation route (Figure 1). The energy generation reaction is catalyzed by a heterologously expressed formate dehydrogenase from *Pseudomonas sp.* strain 101 (PsFDH), which oxidizes formate to CO_2_ to regenerate NADH [17]. The formate assimilation route was subdivided into three modules to facilitate stepwise integration and confirmation of individual modules in *E. coli*. In the first (C1) module, formate is ligated to tetrahydrofolate (THF) at the expense of ATP and reduced to 5,10-methylene-THF by three heterologously expressed enzymes originating from *Methylorubrum extorquens* (formerly *Methylobacterium extorquens*). Next, in the C2 module the C1 moiety of 5,10-methylene-THF is condensed with CO_2_ and NH_3_ resulting in the two-carbon amino acid glycine. This reaction is catalyzed by the (overexpressed) native glycine cleavage system (Gcv) of *E. coli*, operating in the reverse (reductive) direction, driven in this direction by elevated CO_2_ concentrations.

**Figure 1.**
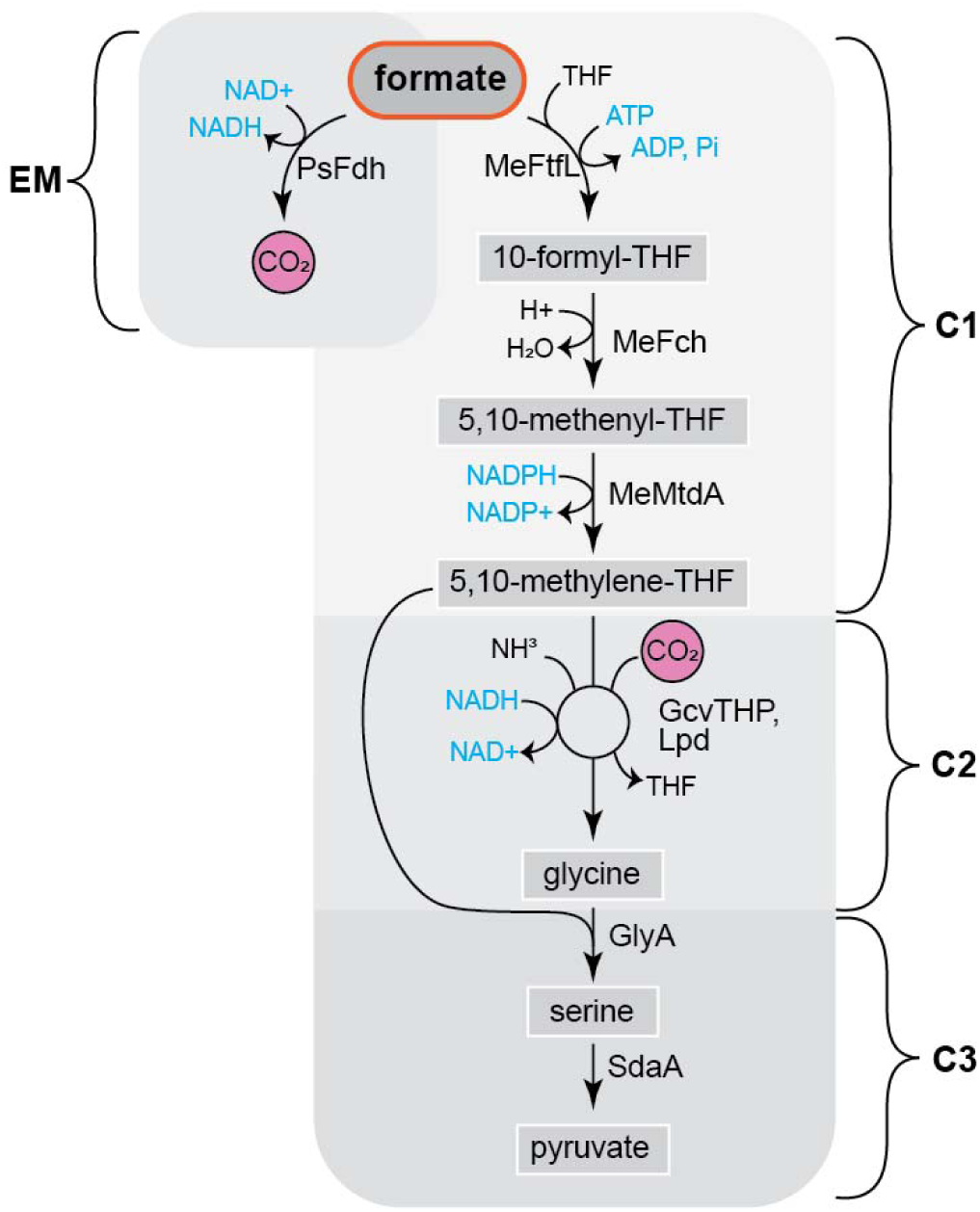
Reductive glycine pathway (rGlyP) as engineered in E. coli by Kim et al. [16]. Me: M. extorquens, PsFDH: formate dehydrogenase, MeFtfL: formate-THF ligase, MeFch: 5,10-methenyl-THF cyclohydrolase, MeMtdA: 5,10-methylene-THF dehydrogenase, GcvTHP,Lpd: glycine cleavage system, GlyA: serine hydroxymethyltransferase, SdaA: serine deaminase.

Finally, in the C3 module, glycine is ligated with another C1 moiety from 5,10-methylene-THF and converted to the C3 amino acid serine and then into pyruvate, by the native serine hydroxymethyltransferase (GlyA) and serine deaminase (SdaA), respectively. Pyruvate can then be converted to various biomass components via native metabolic pathways and possibly also into various bioproducts.

In 2020, Kim et al. [16], achieved full formatotrophic growth of *E. coli* by introducing the complete rGlyP into its genome and overexpressing the native enzymes involved in the rGlyP. This initial strain was named K4 and had a doubling time (DT) of 65-80 h on 28 mM of formate. K4 served as a base strain for further engineering to investigate and improve formatotrophic growth. Supplementary Table 1 gives an overview of growth rates of the various formatotrophic *E. coli* strains derived from K4.

Adaptive laboratory evolution (ALE) was applied to the K4 strain by Kim et al. [16], resulting in the K4e strain, which grew at a DT of 7.7 h on 153 mM formate. Thus, ALE achieved here not only an increased growth rate but also increased formate tolerance as the parental strain K4 did not grow at formate concentrations above ∼60 mM. Two key mutations were further investigated for involvement in this vast growth improvement. These mutations appeared to increase expression of the NADH-regenerating PsFDH, and the native membrane bound transhydrogenase complex (PntAB), which regenerates NADPH by oxidizing NADH. These two mutations were reverse engineered in the naive K4 strain, which did improve growth rate and formate tolerance substantially, but not completely to the extent of the K4e. Unlike K4e, the reconstructed strain (K4 g-FDH* g-PntAB*) did not grow on 153 mM formate. On 109 mM formate the reconstructed strain had a DT of 9.7 h, which is slightly slower than the 7.9 h DT of K4e at the same formate concentration.

After further ALE on K4e, Kim et al. [18] isolated four variants of an even further evolved strain (K4e2), which had a DT of only 6 h on 150 mM formate and a further improved formate tolerance, as well as an increased biomass yield compared to K4e from 2.3 to 3.3 g_CDW_ mol^-1^ formate. The growth improvement of K4e2 was in part attributed to decreased acetate production due to a mobile genetic element integration interrupting the promotor region of the gene encoding acetate kinase (*ackA*).

Acetate kinase and phosphate acetyltransferase (Pta) together catalyze the conversion of acetyl-CoA to acetate. During fast growth on glucose, this conversion is used as part of overflow metabolism to allow faster growth by investing less resources into respiratory chain enzymes while still yielding some energy. However, acetate production during growth on formate via the rGlyP is unfavorable as it has a net energy cost and does not contribute to biomass formation. Furthermore, the potential re-uptake of acetate costs additional ATP. Hence, Kim et al. [18], subsequently deleted the full *ackA-pta* operon in K4e2 as a more stable and rational approach to decrease acetate production, which resulted in slightly better growth than K4e2 with the interrupted *ackA* promoter. This *ackA-pta* deletion was also reconstructed in the predecessor strain K4e. However, while the reconstructed strain K4e Δ*ackA-pta* had a decreased DT (from 8.5 to 7.2 h on 150 mM formate), it did not grow as fast as the fully evolved K4e2 Δ*ackA-pta* (6.4 h). Thus, it appears that blocking acetate production through the AckA-Pta route does not explain the full extent of the K4e2 improvement over K4e. Other mutations in all four isolates of K4e2 were found in genes encoding the pyruvate dehydrogenase complex regulator (PdhR) and RNA polymerase subunit β’ (RpoC) [18].

In a parallel effort, full formatotrophic growth of *E. coli* via the rGlyP was also demonstrated in another study, however, the best strain resulting from this study (FC8) had a maximum growth rate of only ∼65 hours [19].

To summarize, while some mutations have been demonstrated to contribute to the growth improvement of the evolved strains K4e and K4e2 over their parental strain K4, there should be other mutations also contributing to the improved growth phenotypes of the evolved strains. In this study we further investigated the *E. coli* K4, K4e, and K4e2 strains at a genomic and proteomic level to further identify mutations and their effects on protein levels, aiming to further understand how *E. coli* evolved towards more optimal formatotrophic growth.

The K4e and K4e2 strains have recently also been used as proof-of-principle studies to make relevant bioproducts lactate and polyhydrobutyrate from formate [18], [20]. However, the resulting production levels and rates are much too low for economically and sustainable bioproduction from formate. Furthermore, while the formatotrophic growth of K4e2 is much improved over the parental K4 strain, the growth rate and biomass yield are still relatively low compared to natural formatotrophs [6], [21]. This is indicates that the formatotrophic *E. coli* strains still have inefficiencies, such as bottlenecks or sub optimal fluxes in their metabolism, that limit biomass yields and productivities. Therefore, we combined in this work the omics-based approaches with metabolic modelling of the *E. coli* rGlyP metabolism to discover bottlenecks and potential targets for further optimization of these strains for improved growth on formate.

## 2. Methods

### 2.1. Strains and culture conditions

The formatotrophic *E. coli* strains used in this study are listed in Table 1. Lysogeny Broth (LB) medium (10 g/L tryptone, 5 g/L yeast extract, 10 g/L NaCl) was used for rich medium cultivation. M9 minimal medium (50 mM Na_2_HPO_4_, 20 mM KH_2_PO_4_, 1 mM NaCl, 20 mM NH_4_Cl, 2 mM MgSO_4_ and 100 μM CaCl_2_, 134 μM EDTA, 13 μM FeCl_3_ ·6H_2_O, 6.2 μM ZnCl_2_, 0.76 μM CuCl_2_ ·2H_2_O, 0.42 μM CoCl_2_ ·2H_2_O, 1.62 μM H_3_BO_3_, 0.081 μM MnCl_2_ ·4H_2_O, set to pH 7.4) was used for all cultivations on formate. All cultivations were carried out at 37⁰C, with orbital shaking at 180 rpm. All cultivations on formate were carried out in incubators with 10% CO_2_ and 90% air.

**Table 1.**
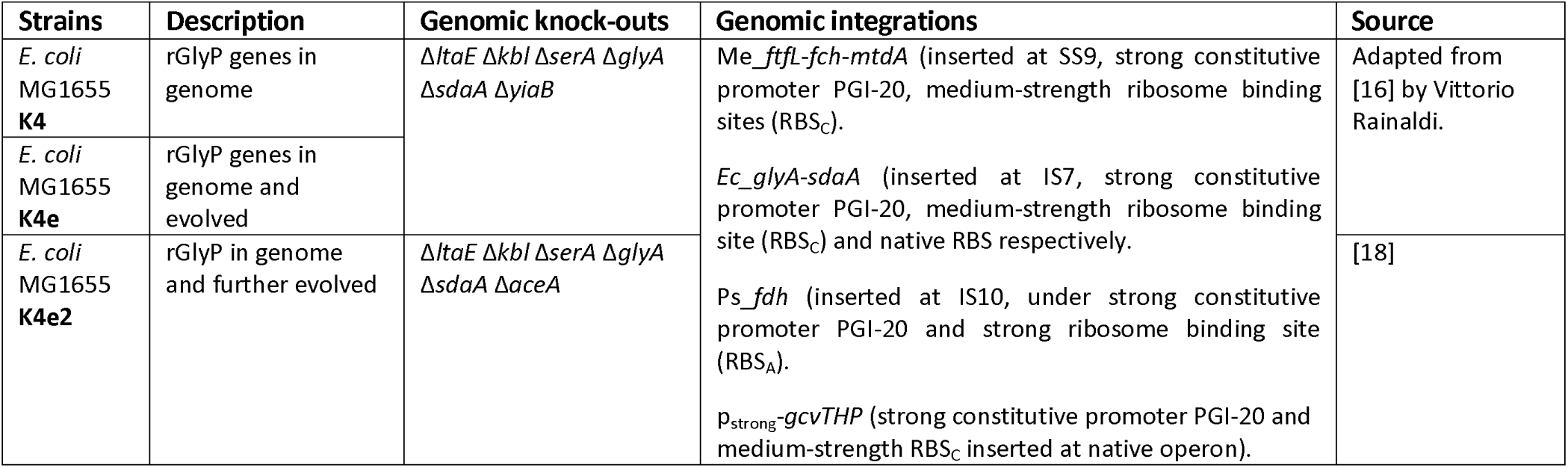
E. coli strains used in this study.

### 2.2. Whole genome sequencing

Cells were grown overnight on LB and DNA extraction was performed using the DNeasy® Blood & Tissue kit manufactured by QIAGEN, according to manufacturer’s instructions (version 07/2020) for Gram-negative bacteria, including the optional RNase A digestion step. Whole genome sequencing was performed by Novogene using the Illumina NovaSeq 6000 platform for paired-end reads of 150bp. Genome sequence data was analyzed using Geneious 10.0.9. Paired reads were mapped to a refence genome (*Escherichia coli* str. K-12 substr. MG1655, <u>U00096.3</u>) retrieved from NCBI using default settings for mapping to reference genome, and a consensus sequence was generated based on majority of reads.

### 2.3. Quantitative proteomics

#### Cultivation and harvesting

Cell material for proteomics analysis was obtained from cultivations in 10 mL M9 with 30 mM (K4) or 80 mM (K4e & K4e2) formate, inoculated from pre-cultures on the same medium. These cultures were grown in 125 mL vented and baffled Erlenmeyer flasks and harvested during log growth phase (harvesting OD_600_ K4: 0.16-0.18, K4e: 0.54-0.67, K4e2: 0.30-0.43). The equivalent of 3 mL of OD_600_=1 was harvested (e.g. 6mL of OD_600_=0.5). These cells were harvested by centrifugation (4500g, 10min, 4⁰C), and washed two times with 1 mL phosphate buffered saline (PBS) (8 g/L NaCl, 0.2 g/L KCl, 1.42 g/L Na_2_HPO_4_, 0.24 g/L KH_2_PO_4_, pH 7.4), then centrifuged again and supernatant was removed. Pellets were flash frozen in liquid nitrogen and stored at -80⁰ C.

#### Protein extraction and alkylation

Cell pellets were resuspended in 300 µL lysis buffer (2% sodium-lauroyl sarcosinate in 100 mM ammonium bicarbonate) and heated for 10 minutes at 90⁰C, shaking. The lysate was further sonicated by 60 ultrasonic pulses (VialTweeter, Hielscher Ultrasonics) and then samples were centrifuged for 10 min at 14,000 rpm at 4°C. Protein concentrations were determined using the Pierce™ BCA Protein Assay Kit (ThermoFisher Scientific), according to manufacturer’s instructions. Then, 7.5 µL of TCEP buffer (0.2 M Tris (2-carboxy-ethyl) phosphine in 100 mM ammonium bicarbonate) was added and incubated for 10 min at 90⁰C, shaking. Next, 7.5 µL of 0.4 M iodoacetamide was added and samples were incubated for 30 min at 25⁰C, shaking (shielded from light).

#### Protein digestion

Based on the BCA results, 50 µg of total protein was digested by addition of 1 µg trypsin (Serva) in diluted lysis buffer (0.25% sodium-lauroyl sarcosinate with 100 mM ammonium bicarbonate) and overnight incubation at 30⁰C. After digestion, SLS was precipitated by adding a final concentration of 1.5% trifluoroacetic acid (TFA, Thermo Fischer Scientific). Peptides were desalted by using C18 solid phase extraction cartridges (Macherey-Nagel). Cartridges were prepared by adding acetonitrile (ACN), followed by equilibration with 0.1% TFA. Peptides were loaded on equilibrated cartridges, washed with 5% ACN and 0.1% TFA containing buffer and finally eluted with 50% ACN and 0.1% TFA.

#### LC-MS

Dried peptides were reconstituted in 0.1% trifluoroacetic acid and then analyzed using liquid chromatography mass spectrometry (LC-MS) carried out on an Exploris 480 instrument connected to an Ultimate 3000 RSLC nano and a nanospray flex ion source (all Thermo Scientific). Peptide separation was performed on a reverse phase HPLC column (75 μm x 42 cm) packed in-house with C18 resin (2.4 μm; Dr. Maisch). The following separating gradient was used: 94% solvent A (0.15% formic acid) and 6% solvent B (99.85% acetonitrile, 0.15% formic acid) to 25% solvent B over 40 minutes, and an additional increase of solvent B to 35% for 20min at a flow rate of 300 nl/min.

MS raw data was acquired on an Exploris 480 (Thermo Scientific) in data independent acquisition (DIA) mode. Peptides were ionized at a spray voltage of 2.3 kV, ion transfer tube temperature set at 275 °C, 445.12003 m/z was used as internal calibrant. The funnel RF level was set to 40. For DIA experiments full MS resolutions were set to 120.000 at m/z 200 and full MS, AGC (Automatic Gain Control) target was 300% with an IT of 50 ms. Mass range was set to 350–1400. AGC target value for fragment spectra was set at 3000%. 45 windows of 14 Da were used with an overlap of 1 Da. Resolution was set to 15,000 and IT to 22 ms. Stepped HCD collision energy of 25, 27.5, 30 % was used. MS1 data was acquired in profile, MS2 DIA data in centroid mode.

#### Data analysis

DIA data was analyzed using DIA-NN [22] as previously described by [23] but using the Uniprot database for *E. coli* with the following additional protein sequences: Formate-THF ligase, 5,10-methenyl-THF cyclohydrolase, and 5,10-methylene-THF dehydrogenase from *M. extorquens AM1*, and formate dehydrogenase from *Pseudomonas* sp. (strain 101). Each with an N-terminal His-tag (complete sequences in Supplementary). Full tryptic digest with up to three missed cleavage sites, oxidized methionines, and carbamidomethylated cysteines was set, with match between runs and remove likely interferences enabled. The neural network classifier was set to the single-pass mode, and protein inference was based on genes. Quantification strategy was set to any LC (high accuracy). Cross-run normalization was set to RT-dependent. Library generation was set to smart profiling. DIA-NN results were further processed using SafeQuant. DIA-NN outputs were further evaluated using the SafeQuant script modified to process DIA-NN outputs [24], [25].

The mass spectrometry proteomics data have been deposited to the ProteomeXchange Consortium via the PRIDE partner repository with the dataset identifier PXD050315 and in Supplementary data 1.

### 2.4. Constraint-based metabolic modelling

#### Model

The iJO1366 genome-scale metabolic model of *E. coli* MG1655 developed by Orth et al. [26] was modified and used in this study. Modifications to the model included adding the reactions in the rGlyP pathway not present yet in the model. For this, an NADH-regenerating formate dehydrogenase reaction (ID: FDH) was added and the glycine cleavage system reaction (ID: GLYCL) was made reversible. Additionally, the lower bound of the ATP maintenance reaction (ID: ATPM) was updated from 3.15 to 6.86 mmol g_CDW_ ^−1^ h ^−1^ to match the most recent *E. coli* MG1655 model (iML1515) [27] and the NAD(P) transhydrogenase (periplasm) reaction (ID: THD2pp) was updated to import only one proton into the cytoplasm instead of two [28], [29]. Furthermore, the NADP-dependent malic enzyme reaction (ID: ME2) was made reversible and the malate oxidase reaction (ID: MOX) was made irreversible (only allowing conversion of L-malate to oxaloacetate).

#### pFBA and Flux sampling

For each growth condition the optimal value of the selected objective function (either growth or production of target metabolites) was predicted using the flux balance analysis (FBA) tool from the COnstraint-Based Reconstruction and Analysis package for python (COBRApy) [30].

Carbon source uptake rates for both growth on formate and growth on glucose were fixed to correspond to the actual growth rate of the fastest growing rGlyP strain K4e2 on formate (namely ∼0.12 h^-1^). To simulate formatotrophic growth, formate uptake rate was set to 22.5 mmol g_CDW_ ^−1^ h ^−1^ and glucose uptake rate to zero. Growth on glucose was simulated with a glucose uptake rate of 1.5 mmol g_CDW_ ^−1^ h ^−1^ and formate uptake rate of zero.

The constraint-based model was adapted to run parsimonious Flux balance Analysis (pFBA) [31]. For this adaptation, reversible reactions were split into two reactions, forward and reverse. In addition an additional ‘flux’ metabolite was added as a product to each reaction and a sink reaction was introduced for this flux metabolite. The FBA objective was set to minimize the flux through the flux metabolite sink. This gave the minimal amount of flux the model needed to still produce the optimal growth rate. Then, the flux through the flux metabolite sink was fixed to the minimal number allowing a small deviation of 0.01 mmol h ^−1^. After this flux sampling was performed; 50000 samples were taken for each substrate using the optGp methods [32].

From the flux samples the following statistics were calculated for each reaction: average, standard deviation and the Shapiro– Wilk test statistic for normal distribution [33]. Prior to the calculations, fluxes through reverse reactions were subtracted from fluxes through the corresponding forward reaction. For the relevant comparisons the Kolmogorov–Smirnov test statistic and log2 fold change statistical calculations were done using the python SciPy library

#### Calculating the cost of pathways and energy cofactors

The efficiency of various pathways was calculated by adding sink reactions for relevant metabolites (in the cytoplasm) and running FBA setting the flux through the sink reaction as maximization objective. Pathway efficiencies were tested by eliminating competing reactions as indicated. For a fair comparison of different pathways, NADH was always produced via NADH-regenerating formate dehydrogenase, NADPH via NAD(P) transhydrogenase and ATP via NADH dehydrogenase (ubiquinone-8 & 3 protons), cytochrome oxidase bo3 (ubiquinol-8: 4 protons) and ATP synthase (four protons for one ATP), as the full model predicts that this is the way most of these energy cofactors are made when growing on formate. In addition, pathway efficiency was computed by setting maintenance requirements to zero. When calculating the cost of individual metabolites, we confirmed that the given metabolite had a net cost of ATP, NADH, and NADPH. While growth on formate virtually always resulted in a net cost of these energy carriers, growth on glucose for instance could for some metabolites result in a net yield of one energy carrier and a net cost of another, leading to conversions of energy carriers that would not be necessary (to the same degree) in full biomass production.

To estimate the potential to generate ATP, NADH or NADPH conversion reactions were introduced converting each of them to their low energy form (ADP, NAD^+^ and NADP^+^, respectively). The corresponding reaction was set as FBA maximization objective in each case. By dividing the maximal flux by the formate input flux, the cost/formate was obtained.

## 3. Results & Discussion

We investigated three *E. coli* strains (K4, K4e, and K4e2), which were engineered and evolved to grow on formate through the synthetic rGlyP [16], [18]. We aimed to characterize the metabolic network of these strains and identify targets for future optimization, through a combination of omics-based data collection and metabolic modelling.

The main product of the rGlyP is pyruvate, which can then be converted through central carbon metabolism further into 11 other biomass precursors and from there to the rest of biomass [34]. We have predicted the most efficient metabolic routes for production of these main precursors from formate, using a genome-scale model of *E. coli* metabolism adapted from Orth et al. [26] to include the rGlyP pathway. We also compared the predicted metabolic fluxes for optimal growth on formate for *E. coli* via the rGlyP with predicted fluxes for growth on its natural substrate glucose (Figure 2). In parallel, we analyzed the proteome of *E. coli* strains K4, K4e, and K4e2 during growth on formate, and compared these to previously published proteomes for *E. coli* grown on glucose at a controlled growth rate, similar to the growth of the fastest formatotrophic strain K4e2 on formate (∼6 h doubling time) (Figure 3). We compare protein allocation and the optimal flux network for the growth on formate, with those of the ancestor strain when growing on glucose, assuming that *E. coli* is better optimized during its long evolutionary history for the natural substrate. From this comparison we aim to identify enzymes and related reactions that are possibly not optimally adapted yet for growth on formate.

**Figure 2.**
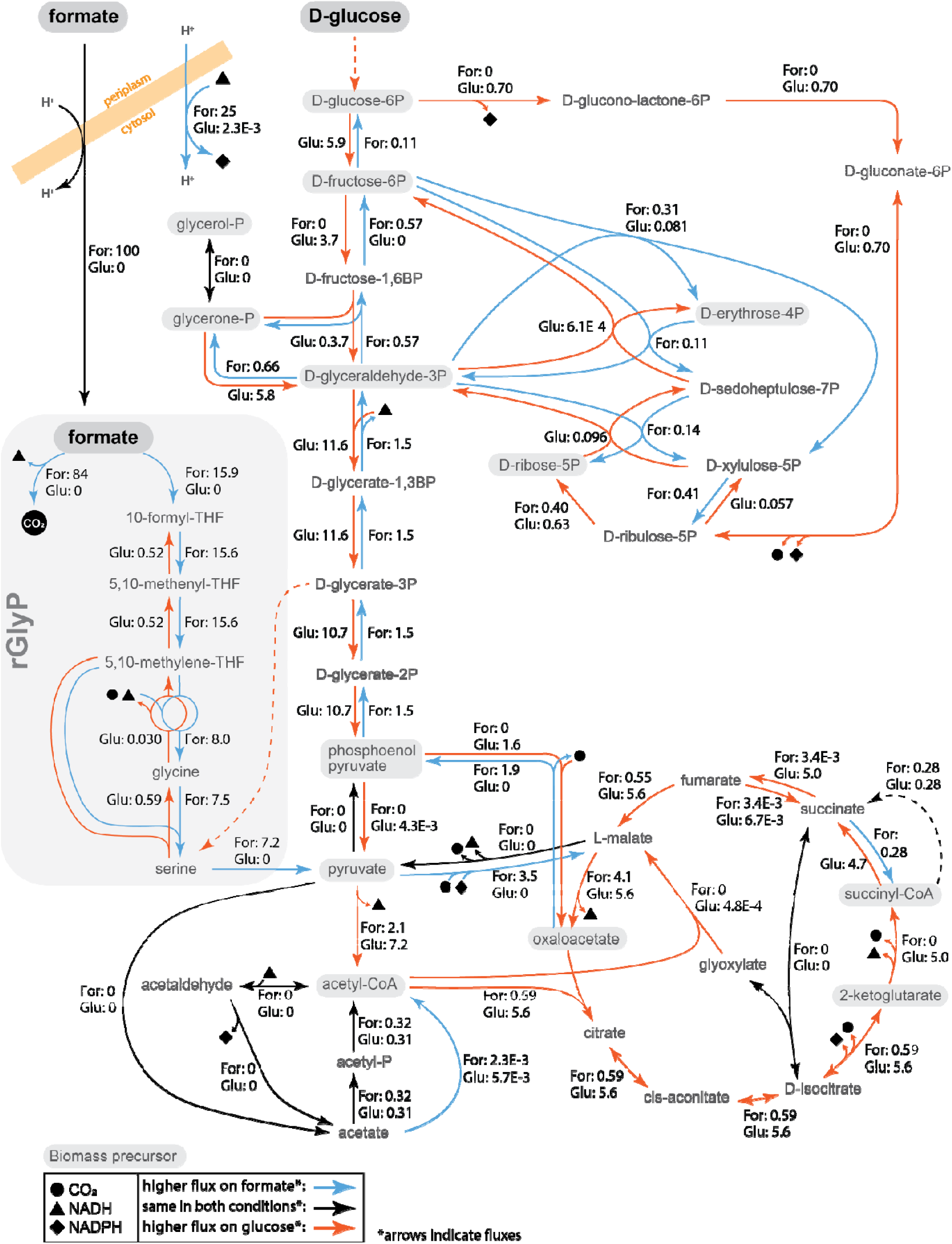
Average predicted fluxes using the metabolic model for both growth on formate and glucose at the same growth rate (∼6h doubling time), all fluxes are normalized for formate uptake flux of 100. Blue arrows denote reactions that, based on the model, carry a higher flux for formatotrophic growth than for growth on glucose, while orange arrows indicate reactions with a higher flux on glucose. Reactions with both blue and orange arrows have a reversed flux direction between two conditions. Black arrows indicate fluxes that are unchanged across the

**Figure 3.**
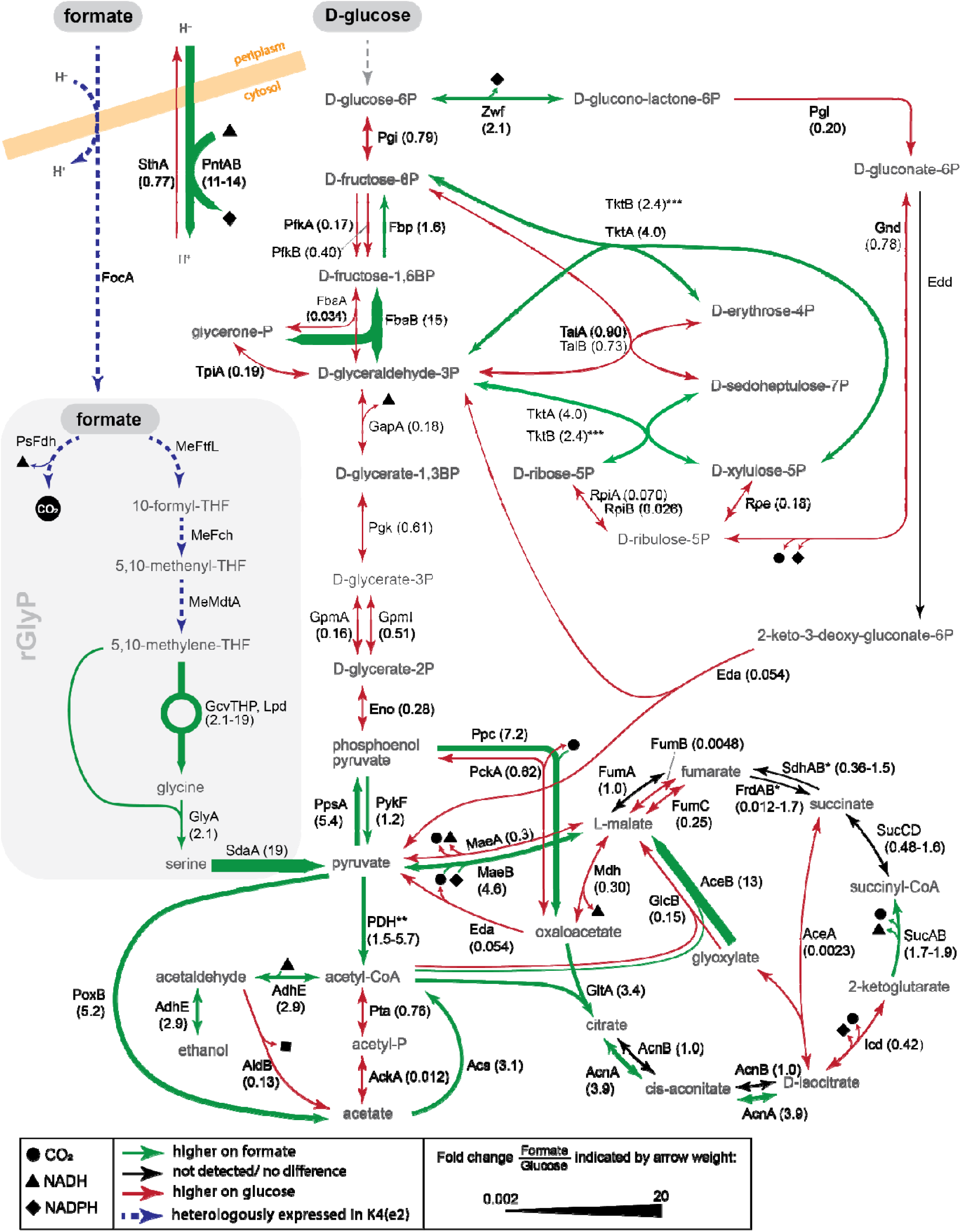
Protein abundance fold changes for the formatotrophic grown E. coli K4e2, versus E. coli grown on glucose. Green arrows denote protein abundance is higher on formate than on glucose, red arrow denote the opposite (higher on glucose than on formate). Black arrows indicates either no difference, or that a protein was not detected. Arrow thickness indicates degree of fold change (thicker arrow is much more abundant on formate than on glucose, thinner arrow means much less abundant of formate than on glucose)

When analyzing the formatotrophic growth of *E. coli*, the metabolic model predicts a maximum yield of 5.4 g mol^-1^ formate, which is still much higher than the highest achieved experimental yield (3.3 g mol^-1^ formate). However, predicted yields are highly dependent on the non-growth-associated ATP maintenance (6.86 mmol g_CDW_ ^−1^ h ^−1^ assumed in the model), which may change in different growth conditions. Nevertheless, yields of ∼4 g mol^-1^ formate have been reported for some formatotrophs using the CBB cycle, which is less energetically efficient than the rGlyP [21], [35], [36]. Furthermore, recently a yield of 4.5 g mol^-1^ formate was achieved through the rGlyP engineered in *C. necator* by Beau Dronsella (personal communication). This yield gap between these other formatotrophs and the rGlyP in *E. coli* indicates that there are potentially energy-losses or stoichiometrically less efficient routes occurring in the formatotrophic metabolism of the currently available formatotrophic *E. coli* strains. The yield gap may also be partly related to the still not optimal growth rate, as increasing growth rate normally also increases yield [37]. The fastest CBB cycle formatotrophs can typically grow slightly faster (∼3.5-4 hours doubling time) [35], [36], [38] than the fastest growth via the rGlyP so far (6 hours).

Here, we discuss key findings from the proteomics and modelling comparison for growth on formate and glucose. We will discuss possible routes and predicted most optimal routes for the regeneration of energy cofactors and the production of biomass precursors from formate. These routes will be analyzed using proteomics data, which provide an estimate for the abundance of the enzymes involved in these routes and hence, a proxy for the flux through these routes.

### 3.1. NADH regeneration by formate dehydrogenase is optimal

During growth on formate, energy metabolism widely differs from growth on glucose because formate is a much more oxidized, single-carbon substrate. The most efficient and direct way to derive energy from formate is through its oxidation by heterologously introduced formate dehydrogenase (PsFDH), which regenerates NADH. In turn, NADH powers the regeneration of other energy carriers. Alternatively, NADH could be regenerated in the TCA cycle, which costs more than twice as much formate per NADH regenerated (Figure 4 A). The costs of regenerating these energy carriers includes the cost of importing formate into the cytoplasm. The mechanism and energetics of formate import into the cytoplasm are not completely understood, though previous studies have suggested that an additional proton is needed to import formate (CHOO^-^) as neutral formic acid (CHOOH), either through diffusion through the membrane or through the hydrophobic center of the FocA transporter located in the inner membrane [39]. Recently, Vanyan et al. [40] showed that in *E. coli*, import of exogenously added formate into the cytoplasm dissipated the proton motive force (PMF), further supporting the notion that formate is likely symported with one (or more) protons. Therefore, the formate import reaction in the model also imports one proton per imported formate. This is why for instance NADH regeneration by PsFDH costs 1.14 formate per NADH, the 0.14 additional costs reflects the energy costs for transport.

**Figure 4.**
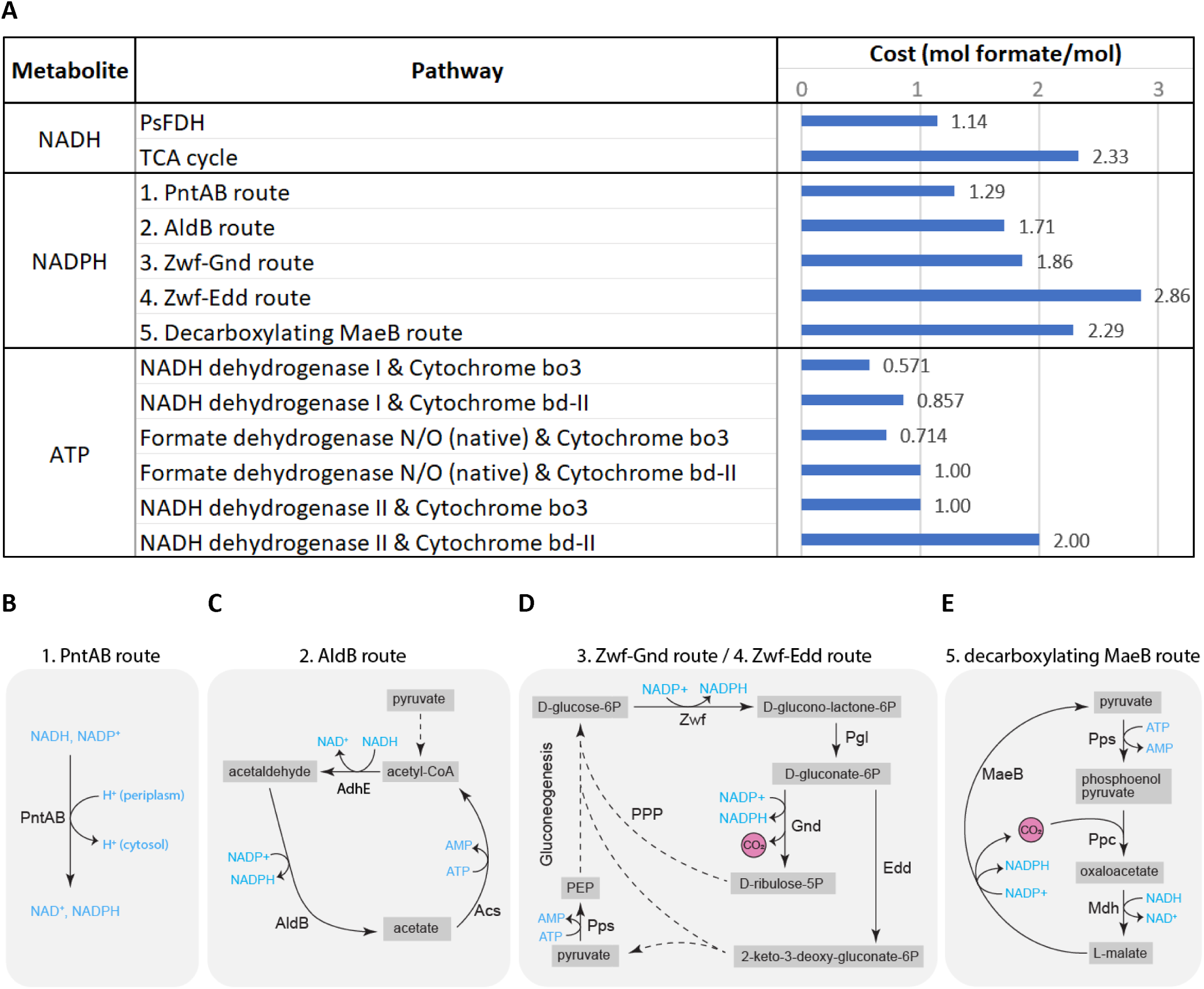
(**A**) The cost of energy carriers NADH, NADPH, and ATP through various regeneration pathways, in mol formate per mol energy carrier. (B-E) Metabolic maps of the five most theoretically efficient NADPH regeneration routes in formatotrophic E. coli strains analyzed by metabolic modelling. Not all reaction components are depicted.

### 3.2. NADPH regeneration by membrane-bound transhydrogenase is optimal

The first part of the rGlyP oxidizes one NADPH to NADP^+^ for each assimilated formate, leading to a high demand for NADPH regeneration. We modelled the top five theoretically most efficient routes for NADPH regeneration that may be used by the formatotrophic *E. coli* strains during growth on formate (Figure 4)).

However, there is a potentially even more efficient route (not shown in figure) through a formate dehydrogenase that directly regenerates NADPH. For instance, the NAD^+^-specific PsFDH enzyme was previously engineered to achieve high kinetic efficiency and specificity for NADPH regeneration. While the strains analyzed in this study do not possess an NADPH-regenerating FDH and thus are unable to perform this route, introducing such an enzyme in would be a promising avenue for future optimization of formatotrophic growth.

The most efficient route currently possible in the formatotrophic strains utilizes membrane bound transhydrogenase (PntAB) to transhydrogenate NADH into NADPH at the cost of one proton translocated into the cell (Figure 4 B). Kim et al. (2020) found mutations in the promotor of the *pntAB* operon, as well as the 5’UTR of the gene encoding PsFDH in the K4e strain, which appeared to increase transcript levels for both. Our proteomics findings also show that the abundance of both these proteins increased, PntAB appeared to be ∼11-13 fold higher in the K4e2 strain on formate than in the glucose reference (Figure 3).

The second most efficient route for NADPH regeneration predicted by the model is a cycle that converts NADH to NADPH via acetaldehyde (Figure 4 C). This cycle starts by first reducing acetyl-CoA with NADH to acetaldehyde and subsequent oxidation to acetate by regeneration of NADPH, catalyzed by two acetaldehyde dehydrogenases (AdhE and AldB respectively). Next, acetate is converted back to acetyl-CoA (costing the equivalent of 2 ATP), by acetyl-CoA synthetase (Acs). This route costs an additional 0.42 mol formate per mol NADPH regenerated compared to the PntAB route (Figure 4 A). We observed increased proteomic abundance of AdhE and Acs on formate compared to glucose, but decreased levels of AldB (Figure 3). Since AldB is the NADPH regenerating reaction, it appears that this route is not being used for NADPH regeneration. According to the model prediction, AdhE should carry no flux (Figure 2), thus may be a suitable target for knock-out.

The next two routes for NADPH regeneration use glucose-6-phosphate dehydrogenase (Zwf) in the (oxidative) Pentose Phosphate Pathway (PPP) to produce gluconate-6-phospate (Figure 4 D). Gluconate-6P can either be oxidized to ribulose-5-phospate to regenerate a second NADPH by 6-phosphogluconate dehydrogenase (Gnd) or it can be dehydrated to 2-dehydro-3-deoxy-D-gluconate-6-phosphate by phosphogluconate dehydratase (Edd). The Zwf-Gnd route and Zwf-Edd route (known as the Entner Doudoroff pathway) respectively use 0.57 mol and 1.57 mol more formate than the PntAB route (Figure 4 A). Hence the model predicts no flux should go through these routes during growth on formate. However, during growth on glucose Zwf-Gnd is one of the key NADPH regeneration routes as also observed in the model simulations (Figure 2).

Interestingly, proteomics indicated a higher abundance of Zwf on formate than on glucose (Figure 3). Notably, the gene encoding Zwf is also known as one of the most consistently flux-coupled genes in *E. coli* [41]. Zwf is regulated by the SoxRS regulon and this regulatory mechanism is controlled by NADPH levels. Low NADPH levels during rGlyP on formate may be a possible explanation for the upregulation of Zwf [42]. Thus, Zwf appears to be a suitable candidate for downregulation or even knock-out, as this upregulation may lead to less efficient NADPH regeneration than via PntAB.

The final NADPH regeneration pathway discussed here is a pyruvate-malate cycle that uses malic enzyme B (MaeB) to decarboxylate malate to pyruvate. Pyruvate is then converted back to malate via phosphoenol pyruvate (PEP) and oxaloacetate, catalyzed by PEP synthase (Pps), PEP carboxylase (Ppc), and malate dehydrogenase (Mdh) (Figure 4 E). This cycle costs one formate more for NADPH regeneration than the PntAB route (Figure 4 A), and requires MaeB to catalyze in its malate-decarboxylating direction. However, it is uncertain which direction MaeB operates *in vivo* during growth on formate, since the conditions during formatotrophic growth (including elevated CO_2_) may push this reversible reaction more towards carboxylation. MaeB levels appear to be higher on formate than on glucose, which could mean that this enzyme is regenerating NADPH, if it does operate in the decarboxylating direction. Alternatively, if MaeB is operating in the carboxylating direction, it may be used for biomass formation, which is further discussed below.

### 3.3. ATP production by NADH dehydrogenase II and cytochrome bo_3_ is optimal

There are various routes for generating the proton motive force (PMF) required for the production of ATP and for other PMF-driven reactions (e.g. NADPH regeneration by membrane-bound transhydrogenase, formate import, etc.). The efficiency of these routes was compared in terms of formate cost per ATP regenerated. By far the stoichiometrically most efficient route for ATP regeneration on formate uses NADH dehydrogenase I (Nuo complex) combined with cytochrome b*o_3_* (Cyo complex) to oxidize NADH and pump protons out of the cytosol (Figure 4 A). Some of the other, less efficient options use combinations with other dehydrogenases and oxidases, namely: NADH dehydrogenase II (Ndh), cytochrome *bd*-I / *bd*-II (Cyd complex / App complex), and formate dehydrogenase N / O (Fdn complex / Fdo complex). Formate dehydrogenase N and O are native membrane proteins that reduce menaquinone, not to be confused with the heterologous NAD^+^-reducing PsFDH.

The model predicted that flux through the Nuo and Cyo complexes should be 1.9 and 1.6-fold higher on formate than on glucose respectively, and that the other routes for PMF generation should carry no flux for optimal biomass yield on formate. The difference in ATP requirement is because during the breakdown of glucose to major biomass precursors there is net ATP production unlike for formate assimilation, which consumes ATP. In addition, the model predicts that ubiquinol is produced in the TCA cycle during growth on glucose but not on formate, which further contributes to the increased flux on formate vs. glucose. However, proteomics showed that NuoBCEFGI had on average 4 fold higher levels in *E. coli* on glucose than in K4e2 of formate, while the other subunits (NuoAHJKLMN) were not detected in the proteomic analyses (Supplementary Table 3). CyoA was approximately 14-fold higher in the glucose condition than on formate, but CyoBCD were not detected either. The seemingly lower abundances of Nuo and Cyo on formate contradicts what the model predicts as optimal, and may thus be suitable targets for upregulation. However, it may also be that in this case protein abundance does not reflect flux, due to for instance differences in substrate or product concentrations (e.g. NADH:NAD ratio) or post translational modifications. Furthermore, too high activity of these respiratory chain enzymes may lower the NADH:NAD ratio too much, which would likely be unfavorable for the thermodynamic feasibility of glycine production by the glycine cleavage system in the rGlyP. Thus, fine-tuning the expression of the Nuo and Cyo subunits may be needed.

Many of the enzymes involved in the less efficient routes for ATP regeneration were not detected or showed contradicting trends for their different subunits. Therefore, it is unclear whether the other involved enzymes require downregulation or knock-out.

### 3.4. TCA cycle not required for energy generation during formatotrophy, but most enzymes are still abundant

During aerobic growth on glucose and all other natural substrates, the TCA cycle oxidizes acetyl-CoA to regenerate the energy cofactors NADH and NADPH. However, energy generation during growth on formate is most efficiently done directly by PsFDH, while operation of the full TCA cycle is relatively inefficient as mentioned previously. During growth on formate, the TCA cycle is therefore only needed to produce biomass precursors.

The model thus predicts lower flux through almost all TCA cycle enzymes on formate compared to glucose (Figure 2). The TCA cycle section between oxaloacetate and alpha-ketoglutarate still carries flux to fulfil the demand of the precursor alpha-ketoglutarate, which requires about 10 times less flux than for growth on glucose. The section between alpha-ketoglutarate and fumarate is predicted to carry no flux in the oxidative direction during growth on formate. Succinyl-CoA is the remaining (minor) biomass precursor in the TCA cycle. A small amount is needed as a CoA donor for the biosynthesis of the amino acids lysine and methionine. This amino acid synthesis releases succinate again, which can be recycled back to succinyl-CoA in the TCA cycle via succinyl-CoA synthetase (SucCD). Hence, only very low fluxes towards succinate or succinyl-CoA are needed for biosynthesis and to sustain the succinate/succinyl-CoA pool in the cells. The model suggests this flux could come from the reductive direction via fumarate, through fumarate reductase. However, in *E. coli*, fumarate reductase is usually only active under anaerobic conditions.

In the last section of the TCA cycle, malate dehydrogenase (Mdh) carries substantial flux. This is related to its involvement in the stoichiometrically most efficient route to produce oxaloacetate and PEP, as will be explained below.

Contrary to the expectations based on optimal flux distribution for formatotrophic growth, proteomics indicate that several TCA cycle enzymes have higher abundances during growth on formate than during growth on glucose (e.g. citrate synthase: GltA, aconitase: AcnA, and alpha-ketoglutarate dehydrogenase: SucAB) (Figure 3).

Though it is unknown whether these increased levels of GltA, AcnA, and SucAB also lead to increased flux, SucA has been identified as a flux-coupled gene (meaning the qualitative change in abundance was closely linked to the qualitative change in flux) [41]. Since SucAB should carry no flux according to our model prediction, the genes encoding this enzyme complex (sucA-sucB) could be a suitable target for knock-out (or knock-down to allow some succinyl-CoA biosynthesis) to decrease wasteful TCA cycle flux and improve formatotrophic growth efficiency.

Citrate synthase (GltA) is known to be subject to catabolite repression during growth on glucose, which may explain why its expression is higher on formate than on glucose. Since flux through GltA should be approximately 10-fold lower on formate than glucose according to the model, the gene encoding GltA is a potential target for down regulation. Expression tuning of AcnA may be less important if GltA-tuning can limit the flux into the TCA cycle.

### 3.5. Carboxylating malic enzyme most efficient route to oxaloacetate and PEP for growth on formate

Oxaloacetate and alpha-ketoglutarate are two major precursors for the production of amino acids and nucleotides, and respectively support roughly ∼17% and ∼12% of biomass production [34], [43]. Alpha-ketoglutarate is generated from oxaloacetate and acetyl-CoA through a part of the TCA cycle. However, there are various routes to produce oxaloacetate (Figure 5 A).

**Figure 5.**
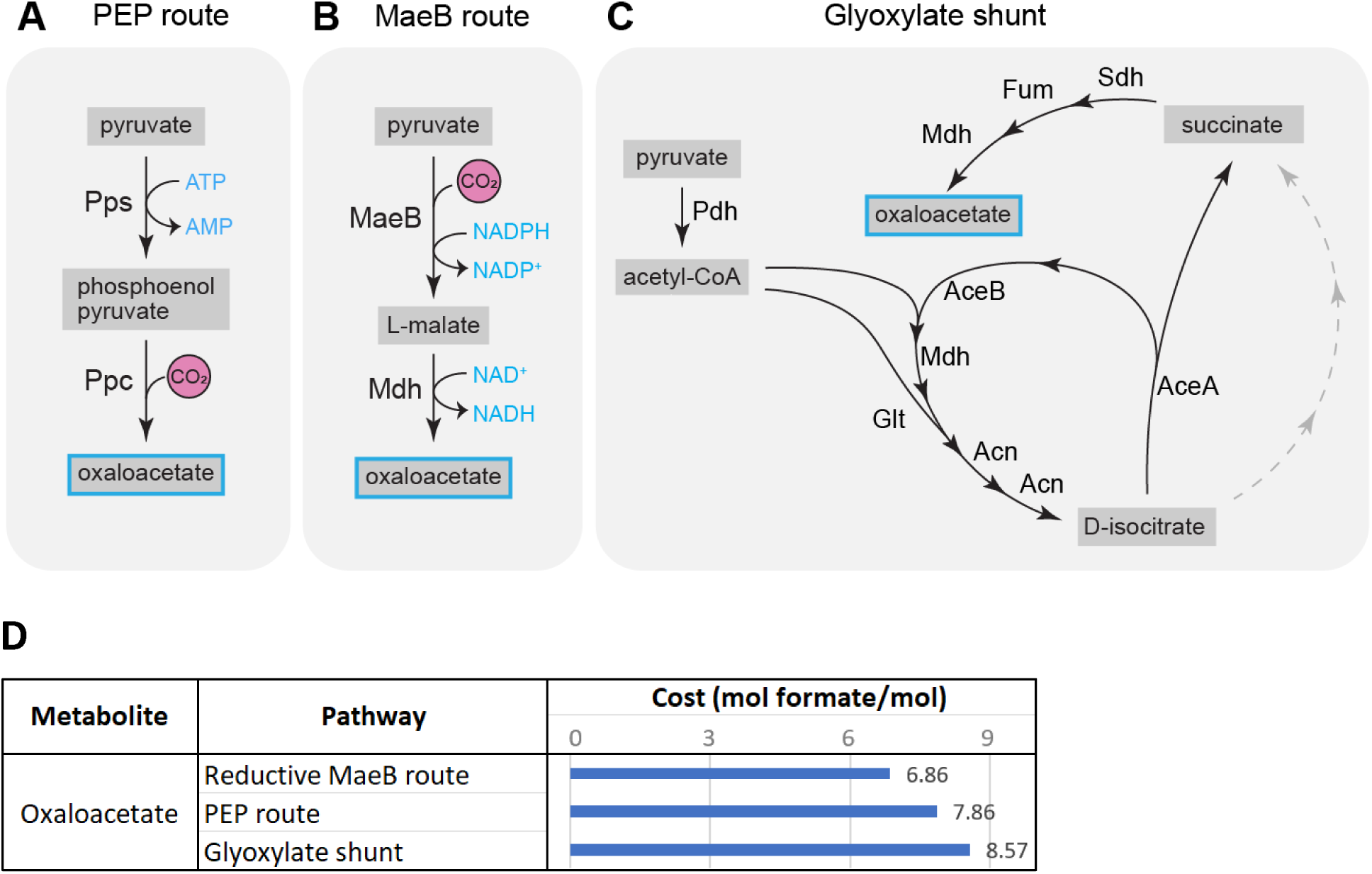
(**A-C**) Metabolic maps of the three oxaloacetate production routes analyzed by metabolic modelling. Pyruvate is produced via the rGlyP as depicted in Figure 1. Not all reaction components are depicted. (**D**) The cost of oxaloacetate biosynthesis through various pathways, in mol formate per mol oxaloacetate.

For growth on glucose, the model predicts that the most efficient route for production of oxaloacetate is through glycolysis until phosphoenolpyruvate (PEP), which can then be converted directly to oxaloacetate by PEP carboxylase (Ppc). For growth on formate, oxaloacetate can also be produced from PEP, which could be generated from pyruvate coming out of the rGlyP via PEP synthase (PpsA) (Figure 5 A). However, the model predicts that oxaloacetate would be generated through a more efficient route, via NADPH-dependent reduction and carboxylation of pyruvate into malate via malic enzyme MaeB (Figure 5 B). This reaction is in the typical *E. coli* model assumed to proceed in the reverse oxidative direction. However, as mentioned previously, this thermodynamically reversible reaction could be functioning in the reductive direction under the conditions for formatotrophic growth including high CO_2_ concentration and possibly elevated NADPH:NADP^+^ (Δ G’^m^ at of 1mM for all reactants is 5 kJ/mol; Δ_r_ G’^m^ at 0.1 atm CO_2_ and 10:1 NADPH:NADP is -3.4 kJ/mol).

Based on the model predictions, the route via PEP synthase costs one additional mol of formate per mol oxaloacetate compared to the route via carboxylating malic enzyme activity (Figure 5 C).

For growth on formate, the route via carboxylating activity of malic enzyme to oxaloacetate is also the most efficient starting route to generate PEP from pyruvate and further precursors that are synthesized via gluconeogenesis from PEP. Hence, the model predicts for growth on formate that all flux towards PEP proceeds from oxaloacetate through PEP carboxykinase (PckA). Relatedly, the model predicts no flux to go via PEP synthetase (PpsA) and PEP carboxylase (Ppc) (Figure 2). However, analysis of the proteome showed that there was an increased level of the latter two enzymes in the *E. coli* strains during growth on formate, whereas the NADPH-dependent enzyme MaeB is also increased (Figure 3). Thus, possibly downregulation of PpsA and Ppc and/or further upregulation of MaeB could potentially increase the yield on formate. For the latter we assume that the reaction through MaeB can indeed proceed fast enough in the carboxylating direction and that the metabolic flux is not yet fully going through that enzyme in the evolved strains. Though in this case there may be a trade-off relationship between efficiency and rate, where the MaeB route could be more stoichiometrically efficient but slower than the more thermodynamically favorable PEP route, potentially resulting in limitations on growth rate.

### 3.6. Glyoxylate shunt is inefficient for biomass production on formate

Another possible route for the production of oxaloacetate from pyruvate is the glyoxylate shunt (Figure 5 C), which can produce oxaloacetate with an additional cost of two mol formate per mol oxaloacetate compared to the MaeB route (Figure 5 D). The glyoxylate shunt is catalyzed by isocitrate lyase (AceA) and malate synthase (AceB/GlcB). The gene encoding AceA has been knocked out during construction of the formatotrophic *E. coli* by Kim et al. [16], and has been restored again in the K4 and K4e strains characterized in this study (for further engineering purposes), but not in K4e2. Thus, since K4e2 lacks AceA, the glyoxylate shunt cannot be used in this strain. However, somewhat surprisingly, proteomics indicate that AceB has a 13-fold higher level in K4e2 on formate than in the glucose reference (Figure 3). Hence, AceB may be a suitable knock-out or downregulation target, to prevent unnecessary proteome allocation to this likely unused enzyme. However, we note that AceB may serve to assimilate glyoxylate, which could be a by-product of an inefficient bypass of the rGlyP. Some of the glycine generated in this pathway may be oxidized to glyoxylate [44], as was also previously observed during rGlyP operation in *Cupriavidus necator* [45]. In the slower growing K4 and K4e strains characterized in this study, where the gene encoding AceA has been reconstituted, proteomic detection of AceA is similar to the glucose reference, though since growth rates between these three conditions vary greatly, a direct comparison cannot be made.

### 3.7. Glycolysis is naturally downregulated in favor of gluconeogenesis during growth on formate

In upper central carbon metabolism, several important biomass precursors are produced. During formatotrophic growth, these metabolites must be produced through gluconeogenesis and the pentose phosphate pathway (PPP).

Glucose-6-phosphate and fructose-6-phosphate are required for the production of glycogen, lipopolysaccharides, and the cell wall. Glycerate-3-phosphate serves as a precursor for glycine, serine, cysteine, and one-carbon metabolites during glycolytic growth. However, the route to produce these compounds from glycerate-3-phosphate was previously knocked out during engineering of the rGlyP in *E. coli*. Since these compounds are also central intermediates in the rGlyP, they are most efficiently directly obtained from this pathway during formatotrophic growth.

As expected, we observe decreased abundance of most glycolytic enzymes during growth on formate, as no major flux is needed, unlike during growth on glucose when glycolysis is a major metabolic pathway (Figure 3). Notable exceptions are enzymes that function specifically in gluconeogenesis and not in glycolysis: FbaB and Fbp. Thus, it appears that gluconeogenesis is already upregulated, though it is unknown whether this is to the optimal level. It could for instance be that gluconeogenesis is too strongly upregulated, as the gluconeogenic demand from the rGlyP is lower than the typical gluconeogenic demand for *E. coli* from other substrates, as compounds generated from glycerate-3-phosphate usually are supplied by the rGlyP directly.

### 3.8. Pentose phosphate pathway products partially made via inefficient Zwf-Gnd route

Erythrose-4-posphate (E4P) and ribose-5-phosphate (R5P) are produced in the PPP and are needed in substantial quantities for synthesis of several amino acids and nucleotides. The most efficient pathway to produce E4P and R5P is direct re-arrangement in the PPP pathway via transketolase (Tkt) and transaldolase (Tal) (Figure 6 A).

**Figure 6.**
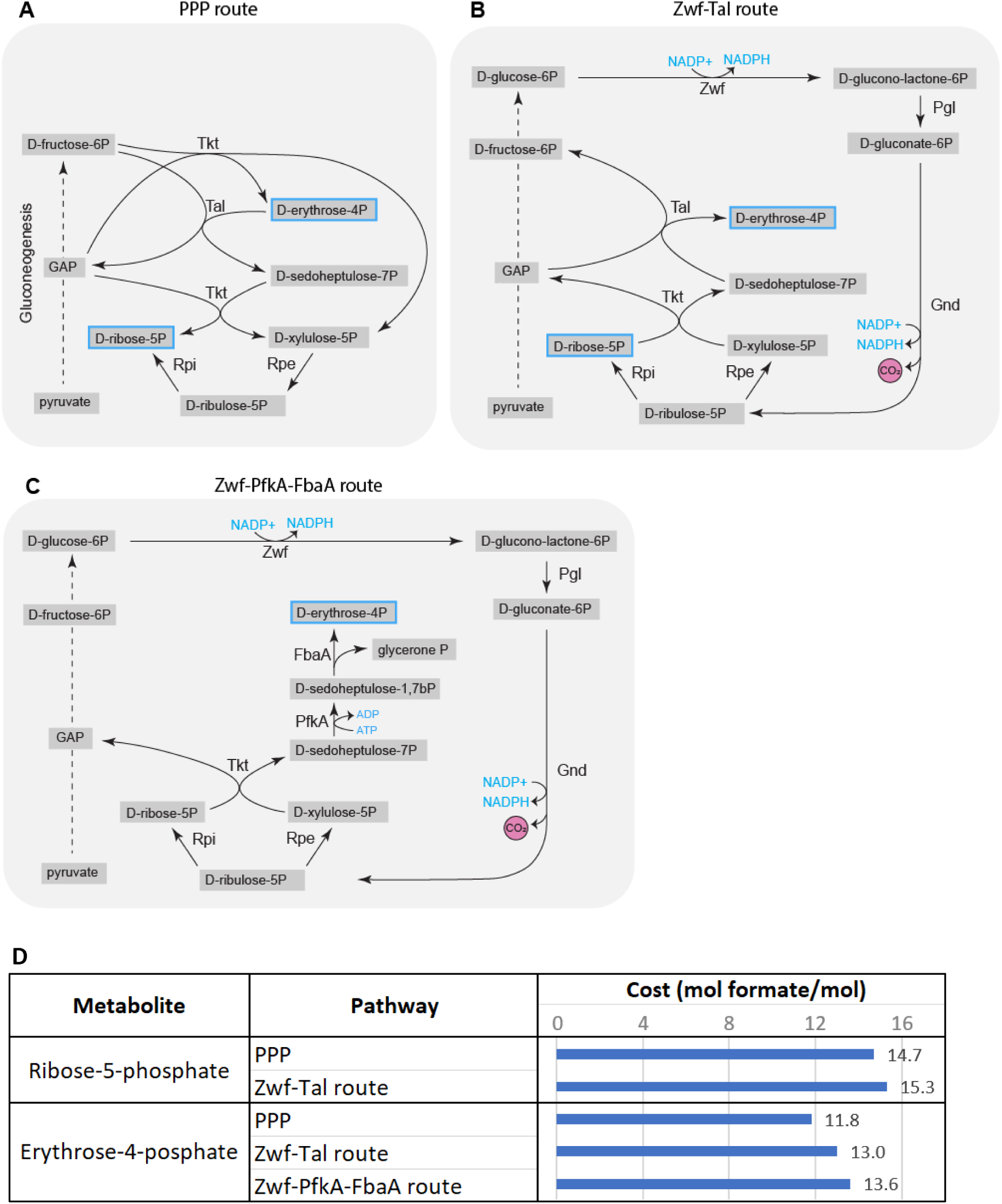
(**A-C**) Metabolic maps of the three Erythrose-4-posphate (E4P) and ribose-5-phosphate (R5P) production routes analyzed by metabolic modelling. Not all reaction components are depicted. (**D**) The cost of oxaloacetate biosynthesis through various pathways, in mol formate per mol oxaloacetate.

Alternative pathways to produce E4P and R5P are via oxidative variants of the PPP utilizing glucose-6-phosphate dehydrogenase (Zwf). The first oxidative PPP variant uses Zwf, as well as Tkt and Tal (Figure 6 B), which costs an additional 1.2 and 0.6 mol formate per E4P and R5P, respectively, compared to the most efficient, non-oxidative PPP route (Figure 6 D). Alternatively, E4P can also be produced via Zwf combined with phosphofructokinase (PfkA) and fructose-bisphosphate aldolase (FbaA) instead of Tal (Figure 6 C), which is costs 1.8 mol formate more than via the PPP (Figure 6 D).

Though the model predicts Tkt, Tal and Rpe should carry more flux on formate than on glucose (Figure 2), from these only TktAB appears to be more abundant at proteomics level on formate (Figure 3). TalAB abundance is similar in both conditions while Rpe appears to be downregulated on formate. Hence, E4P and R5P may be (partially) formed via the less efficient Zwf-Tkt route, as Zwf is more abundant than on glucose, possibly due to the higher NADPH demand in formatotrophic growth. Thus, aiming to redirect the route towards E4P and R5P by downregulating or knocking out Zwf and possibly overexpressed TalAB, and Rpe, may increase metabolic efficiency during growth on formate.

### 3.9. Improved growth of K4e due to improved NAD(P)H regeneration capacity and a potential reduction in flux at rGlyP pathway branchpoint

It has been shown by Kim et al. [16] that the formatotrophic growth improvement of K4e over K4 is largely caused by mutations in regulatory regions of the genes encoding PsFDH and PntAB, the enzymes involved in the most efficient routes for NADH and NADPH generation, respectively. For PntAB the mutation was found in the promotor region, which led to a ∼13-fold increase of transcript level as measured by Kim et al. [16]. This was also corroborated in our findings that protein abundance of PntAB increased by 4.6 and 5.7-fold for PntA and PntB, respectively for K4e compared to K4. The mutation for PsFDH was found at the start of the 5’UTR which caused a 2.5-fold increase in transcript as detected by Kim et al. [16]. We measured a further 13-fold increase of PsFDH at proteome level, which is may be caused by a combination of slightly higher transcript levels as well as improved translation initiation from the mutated 5’UTR.

However, these mutations do not appear to explain the full extent of the K4e phenotype, as their reconstruction in the naïve K4 strain did not yield the same growth rate as K4e.

We further analyzed the genome sequences and identified additional mutations in K4e in the coding sequences of the genes encoding 5,10-methylenetetrahydrofolate reductase (MetF), fatty acid biosynthesis regulator (FabR), and type II secretion system protein (GspL). The mutation in *gspL* is a silent mutation (G138G C->T). This gene is part of an operon that is transcriptionally repressed under standard laboratory conditions [46] and this protein was not detected in any of the proteomes we analyzed. The mutations in *metF* (A280V) and *fabR* (G42V) are amino acid substitutions.

MetF catalyzes the NADH-dependent reduction of 5,10-methylene-THF to 5-methyl-THF, which is part of a native *E. coli* route for methionine synthesis. However, since 5,10-methylene-THF is also an intermediate metabolite in the rGlyP and hence maybe be present in higher concentrations during growth on formate via the rGlyP, MetF expression/activity may need to be downregulated. Proteomics analysis indicates that MetF is indeed somewhat less abundant in K4e than in K4 (Figure 7 A), in addition the amino acid substitution may potentially alter activity further as the substituted amino acid is directly next to an active site residue involved in substrate binding.

**Figure 7.**
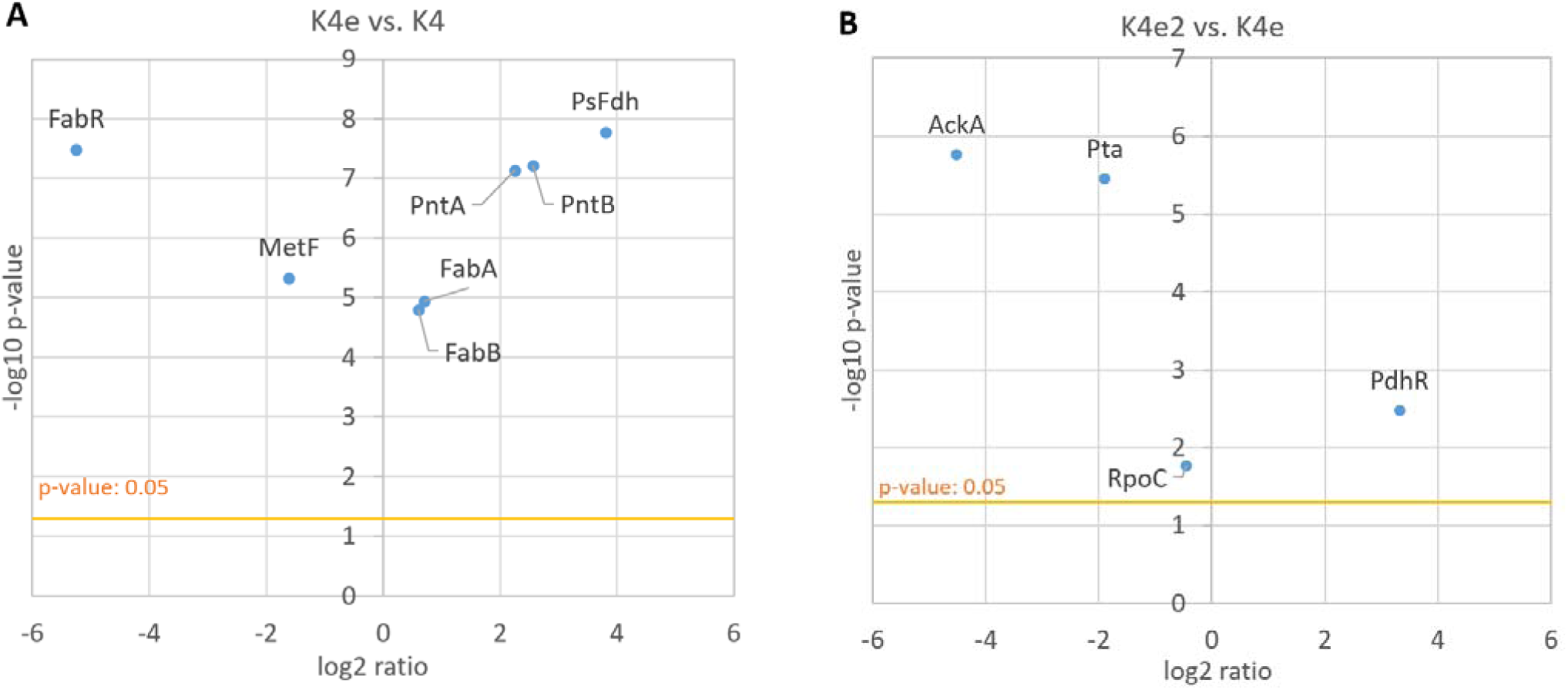
Scatter plots showing abundance differences for proteins encoded by genes that have acquired mutations during ALE. X-axis indicates log2 fold change in protein abundance and y-axis indicates -log10 p-value. (**A**) K4e abundance / K4 abundance, for mutations first occurring in K4e. (**B**) K4e2 abundance / K4e abundance, for mutations first occurring in K4e2

FabR is a transcriptional repressor for the two type II fatty acid synthase enzymes (FabA and FabB), thereby impacting membrane lipid homeostasis [47]. Proteomics analysis indicates that FabR is less abundant in K4e than in K4, and that FabA and FabB are slightly upregulated in K4e (Figure 7 A). However, it is unclear if this upregulation is beneficial for growth on formate and what the mechanistic explanation would be.

### 3.10. Further improved growth of K4e2 due to reduced acetate production, potentially increased Gcv capacity and further proteomic changes

The further growth improvement of K4e2 over K4e was attributed in part to a mutation in the 5’UTR of the *ackA*-*pta* operon, leading to decreased acetate production, as observed by Kim et al. [18]. Proteomics indeed showed a strong downregulation of AckA and Pta in K4e2 compared to K4e (Figure 7 B). In addition, acetate reuptake using Acs also appears to be downregulated in K4e2 compared to K4e (Figure 8 C), though it is not as low as for growth on glucose at the same growth rate (at this growth rate glucose metabolism does not lead to acetate overflow and hence no uptake is required). However, the level of pyruvate oxidase (PoxB) is up in all rGlyP strains compared to glucose. PoxB is a membrane protein coupled to the electron transport chain and converts pyruvate to acetate, which may explain why there is still some production of acetate, even in the K4eΔ*ackA-pta* as Kim et al. [18] reported. Thus, the gene encoding PoxB is likely also a good target for knock-out.

**Figure 8.**
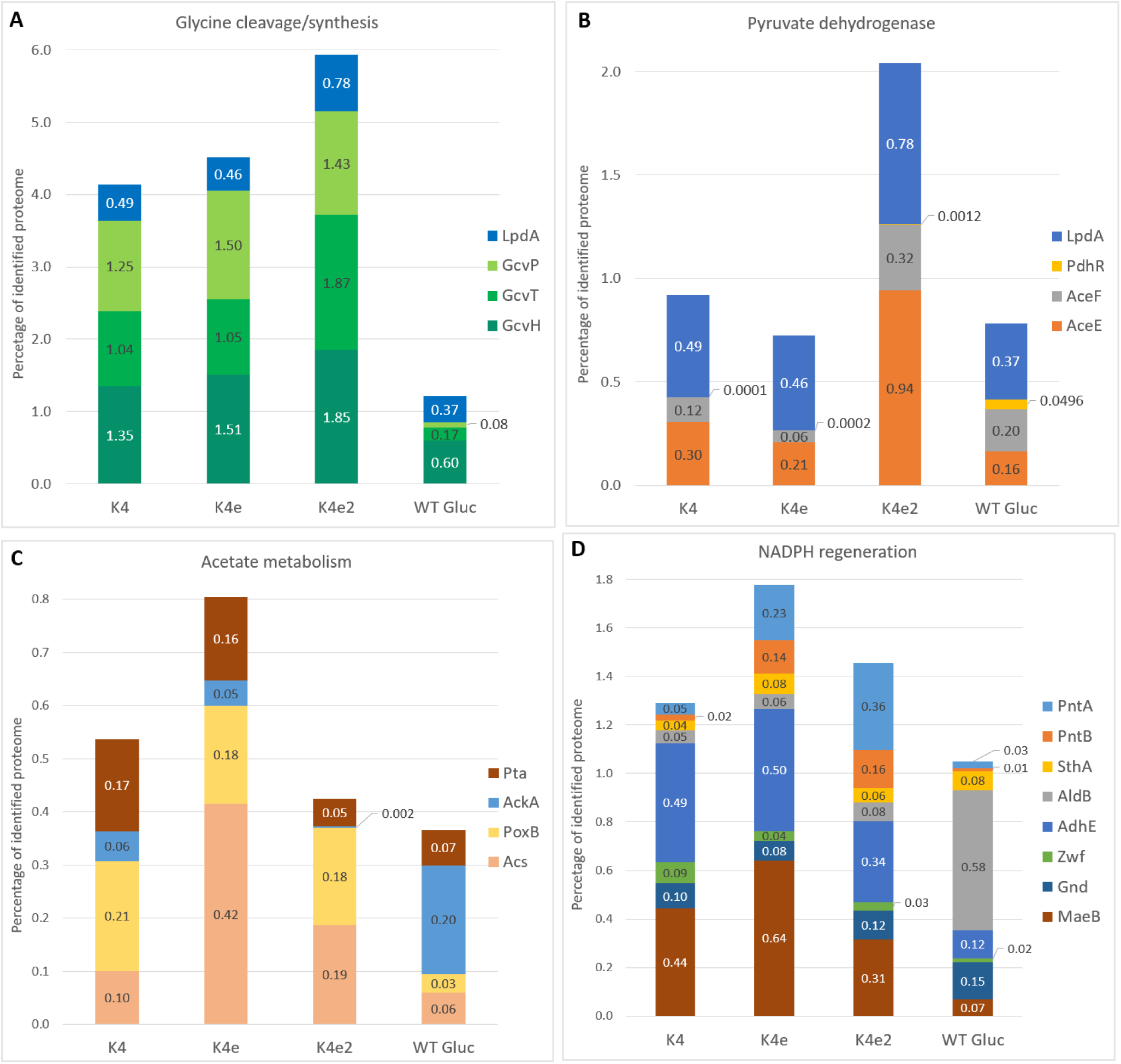
Percentages of total quantified proteome for proteins involved in (**A**) the glycine cleavage system, (**B**) pyruvate dehydrogenase, (**C**) acetate metabolism, (**D**) NADPH regeneration.

Additionally, Kim et al. [18] also identified an amino acid substitution in PdhR and RpoC in K4e2, which likely explains the remainder of the growth differences between K4e2 and K4e.

Since RpoC is a subunit of RNA polymerase involved in DNA binding, it can act as a global regulator of gene expression and is often found mutated in ALE experiments on *E. coli* MG1655 for a range of experimental conditions [48], [49]. While proteomics indicates abundance of RpoC is barely changed in K4e2 compared to K4e (Figure 7 B), the amino acid substitution likely influences RpoC activity. The exact RpoC mutation in K4e2 (A919V) has been reported previously by Szalewska-Palasz et al. [50] to bypass requirement of the alarmone guanosine tetraphosphate (ppGpp) for transcription from sigma-54 promoters. This signaling molecule is involved in the stringent response which is activated during nutrient starvation. ppGpp has many targets, and is for instance involved in decreasing DNA replication and changes in transcriptional profile, inhibiting ribosome maturation and protein translation activity [51]. Possibly, the RpoC mutation could impact transcription levels from starvation sigma-54 promoters compared to the (maintenance associated) sigma-70 promoters. Proteomics showed an increase of proteins expressed from confirmed sigma-54 promoters in K4e2 compared to K4 and K4e, though this could be (partially) caused by differences in growth rate (Supplementary Figure 2). It is known that some sigma-54 genes are associated with nitrogen fixation, though many are not [52]. However, even if the RpoC mutation influences transcription from sigma-54 promoters, it is unclear which (if any) of the impacted genes are contributing to the improved growth of K4e2.

PdhR regulates expression of an operon which encodes the components of the pyruvate dehydrogenase enzyme complex (Pdh), namely pyruvate dehydrogenase E1 component (AceE), pyruvate dehydrogenase E2 subunit (AceF), and lipoamide dehydrogenase (Lpd). We found that levels of PdhR, AceEF, and Lpd are higher in K4e2 compared to K4, K4e, and WT *E. coli* grown on glucose (Figure 8 B), possibly as a result of the PdhR mutation in K4e2. Increased expression of Pdh may not be advantageous since the flux through Pdh should be about three times lower during growth on formate than on glucose, according to the predicted fluxes for optimal yield (Figure 2). However, Lpd is also a component of the glycine cleavage system enzyme complex (Gcv), together with aminomethyltransferase (GcvT), glycine cleavage system H protein (GcvH), and glycine decarboxylase (GcvP). The GcvTHP operon was overexpressed by Kim et al. [16] in K4 to allow growth on formate as the Gcv catalyzes a key reaction in the rGlyP. They did not overexpress Lpd as it was not essential. As expected, proteomics showed increased levels of GcvTHP in the K4 strains on formate compared to WT *E. coli* on glucose (Figure 8 A). We observe that Lpd levels are similar in K4 and K4e on formate, and slightly higher in those strains compared to WT *E. coli* on glucose, but approximately 2-fold higher in K4e2. The upregulation of Lpd in K4e2 may contribute to the strains improved growth on formate by increasing Gcv levels. If this is the case, it may be even better to specifically upregulate Lpd, but not AceEF.

### 3.11. rGlyP strains show trend towards larger proteome fraction of central carbon metabolism

The large differences in growth rates between the K4, K4e, and K4e2 strains on formate likely both result from and cause differences in their overall proteome allocation (Figure 9 A). A notable difference is that the rGlyP fraction of the proteome increased after ALE from approximately 10% in K4, to 13% in K4e, and 16% in K4e2, mainly due to changes in PsFDH and the GCV system. PsFDH abundance largely increased between the K4 and K4e from 0.3% to 3.7% and is even slightly more abundant in K4e2 (4.3%). This increase likely explains a large part of the improved growth between K4 and K4e, as this relatively slow enzyme (k_cat_ ∼10 s^-1^) is responsible for all energy supply in the cell [10], [17]. A further increase of the PsFDH level may boost growth even more, but maybe also burden a large part of the proteome. An alternative may be the use of faster metal-cofactor dependent FDH enzymes, such as the molybdenum-dependent formate dehydrogenase from the natural formatotroph *C. necator.* In another study, the rGlyP was implemented in *C. necator* and in this study it was determined that the CnFdh occupied only ∼1.6% of the proteome to reach doubling times of ∼10 hours [23]. This FDH may also be functionally implemented in formatotrophic *E. coli* strains. Recently, Schulz et al. [53] demonstrated the functional expression of the CnFdh in an *E. coli* NADH sensor strain, which was able to generate all NADH required for *E. coli*. Incorporation of a faster Fdh like CnFDH in K4e2 may be a promising approach to improve growth rate on formate, and an alternative to further increasing overexpression of PsFDH.

**Figure 9.**
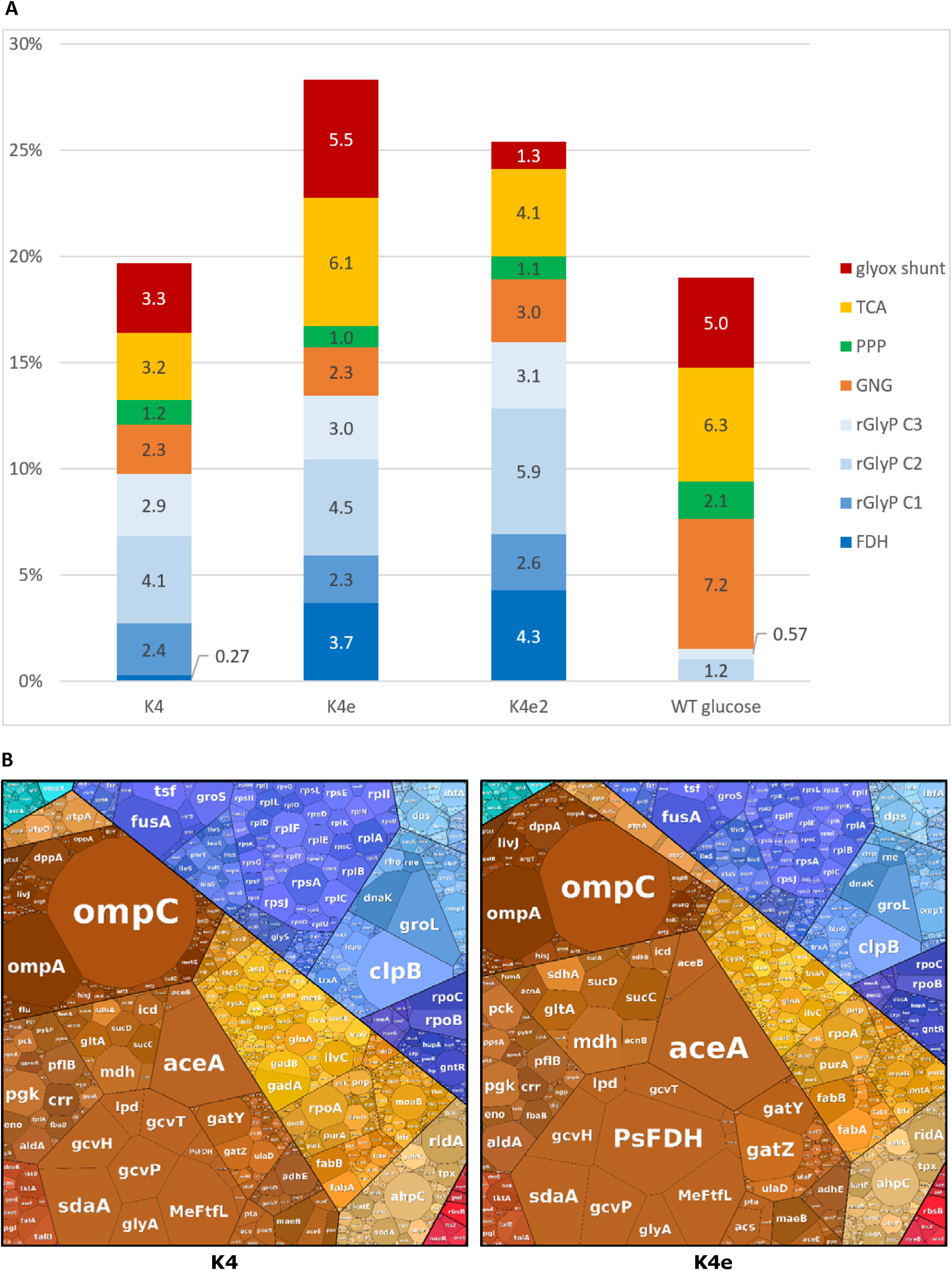
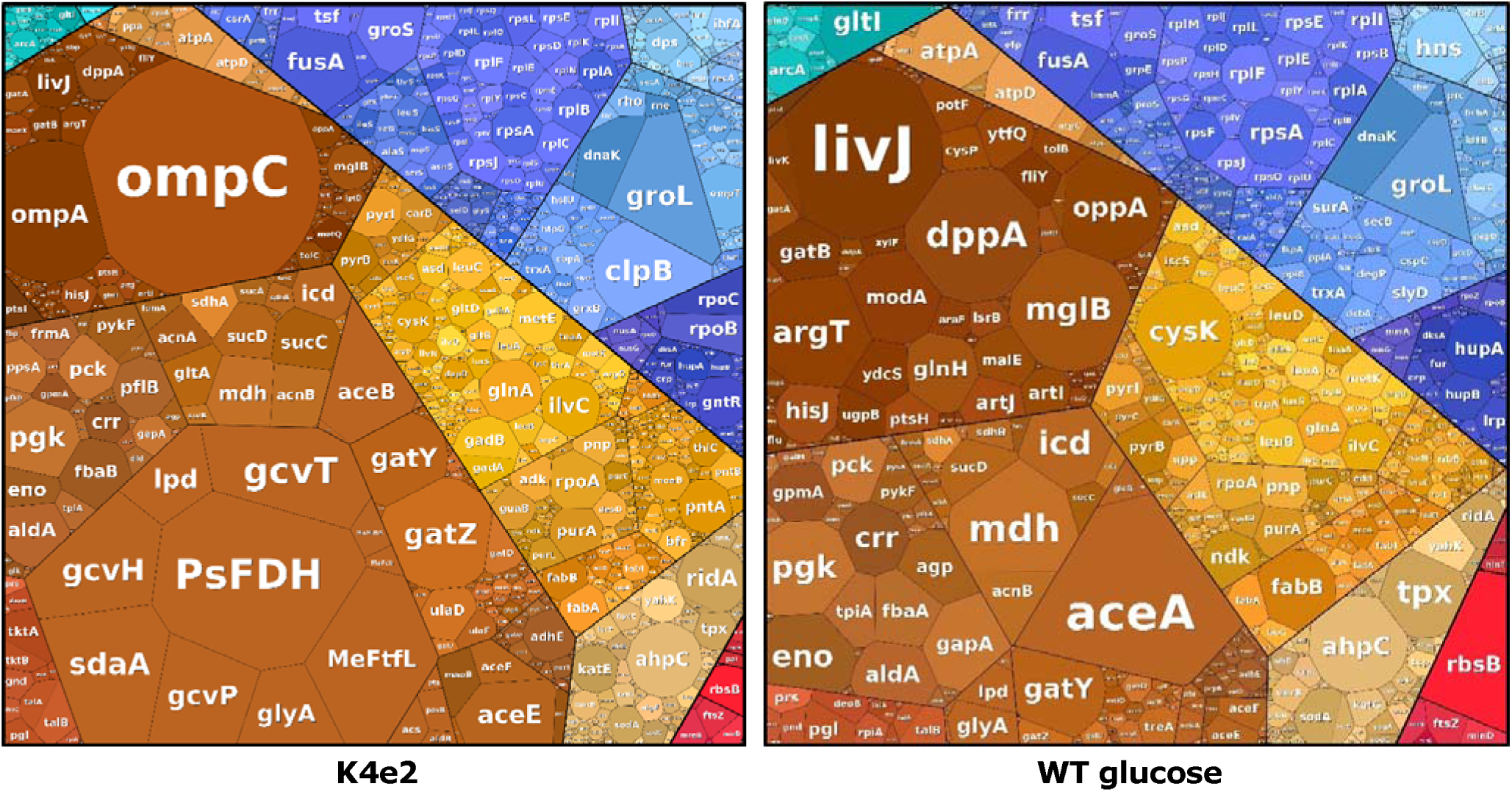
(**A**) Percentages of total quantified proteome for proteins involved in central carbon metabolic pathways. A list of individual proteins included in each pathway is provided in Supplementary Table 2. GNG: gluconeogenesis/ glycolysis. (**B**) Proteomaps displaying all proteins proportional to the size of their proteome fraction, and grouped based on cellular function (Supplementary figure 3). Proteomaps were generated using the tool developed by Liebermeister et al., Nocaj and Brandes, and Otto et al. [61], [62], [63].

The assimilation modules of the rGlyP (C1, C2 and C3) form a relatively stable part of the proteome throughout the different strains. Only between K4e and K4e2, the relative fraction of the C2 module substantially increased from 4.5 to 5.9 %. This is mostly related to the earlier discussed increase in the Lpd enzyme, as well as the GcvT and GcvH proteins that take part in the GCV system. While the kinetics of the GCV system are poorly characterized [54], considering the turnover numbers and the small thermodynamic driving force of this reaction, it is likely that GCV also serves as a bottleneck in the rGlyP pathway. Relatedly, its increased abundance may have contributed to the faster growth rate of K4e2 over K4e.

In total the rGlyP forms a high proportion of the proteome (up to 16%), but during growth on glucose at a similar growth rate to K4e2, the main pathways for carbon assembly and energy generation (glycolysis and TCA cycle) also occupy 13% of the proteome. In addition, several enzymes in the rGlyP have relatively slow kinetics [55]. Furthermore, the rGlyP reactions have to carry higher fluxes of the single-carbon substrate formate, than the fluxes required during the growth on six-carbon substrate glucose in glycolysis and TCA (Figure 2). Thus, an open question remains whether improving the abundance of rGlyP enzymes in the pathway could further boost the growth rate, and which are the bottleneck enzymes in this pathway that would be most essential to further increase.

In the formatotrophic strains, especially in K4e, a large part of the proteome is still taken up by the TCA cycle and glyoxylate shunt (6.1 and 5.5% respectively in K4e), which according to the optimal flux distribution should carry very low or no flux during optimal formatotrophic growth. In K4e2, the fraction of the TCA cycle is already reduced further (to 4.1%) and the glyoxylate shunt occupies a much lower fraction (1.3%), partly because the gene encoding AceA is not present in K4e2.

While the full oxidative TCA cycle does not provide a net biomass yield since it oxidizes the carbon that is fed into it to CO_2_, its activity can be inferred by analyzing the biomass produced from TCA cycle intermediates. Hereto, Kim et al. [16] performed labelling experiments using formate and/or CO_2_ made from the ^13^C isotope of carbon and measured the abundance of ^13^C in several amino acids produced by K4e. In their experiments, threonine (which is produced from oxaloacetate) would be differently labelled depending on whether oxaloacetate was produced through the anapleurotic reactions from pyruvate, or through the TCA cycle via acetyl-CoA. They found that while most threonine was produced through anapleurotic routes, approximately 23% still appeared to come from acetyl-CoA, indicating that the oxidative TCA cycle was still active to some extent. In contrast, Claassens et al. [45] performed similar ^13^C-labeling experiments with the rGlyP in *C. necator*, showing only 12% of threonine came via acetyl-CoA, which may contribute to higher biomass yield of their engineered *C. necator* strain (2.6 g mol^-1^ formate) compared to *E. coli* K4e (2.3 g mol^-1^ formate).

For further optimization of both the protein allocation, and potentially also to reduce wasteful acetyl-CoA oxidation through the TCA cycle, a further reduction of TCA cycle abundance could potentially increase the formatotrophic growth of *E. coli*. Residual TCA cycle flux on the K4e2 may be one of the key wasteful processes that explain the yield gap between the observed and theoretical maximum yield for formatotrophic growth.

In addition to rGlyP proteins, we observed a high abundance of outer membrane porins OmpC and OmpA in the formatotrophic *E. coli* strains (figure 9 B). Both proteins are known to mediate permeability of the outer membrane [56]. They were found *in vitro* to form nonspecific diffusion channels through which various small molecules can passively diffuse, including some sugars and antibiotics, and in the case of OmpC also ions [57], [58], [59], [60]. Thus, these proteins may also facilitate diffusion of formic acid (and perhaps formate) through the outer membrane. However, since all three formatotrophic strains showed this high abundance of OmpC and OmpA compared to the glucose reference, it is unclear whether their expression is upregulated response to growth conditions, or whether this is caused by a genetic difference between the reference and the formatotrophic strains.

## 4. Conclusions

We investigated the metabolism and proteome of the engineered formatotrophic *E. coli* strains K4, K4e, and K4e2 grown on formate, and compared the latter to *E. coli* grown on glucose. We aimed to gain a better understanding of the metabolic differences between these strains and to identify avenues for further improvement of formatotrophic growth by rational engineering. We identified several potential metabolic bottlenecks and inefficient routes in the formatotrophic metabolism of K4e2.

We found that, despite high expression of the heterologous PsFDH that efficiently regenerates NADH, the K4e2 strain may still be using the full TCA cycle for energy generation on formate as well, which is highly inefficient. Thus, improved yield during growth on formate may be achieved by downregulating expression of genes encoding key enzymes in this cycle, such as GltA and SucAB. Furthermore, while the optimal route for NADPH regeneration through PntAB is overexpressed, there may still be a bottleneck for NADPH, possibly contributing to utilization of less efficient NADPH regeneration routes, such as those using Zwf. Thus, knockout of Zwf and stronger overexpression of PntAB may also help improve the metabolic efficiency and biomass yield.

While malate decarboxylation by MaeB is part of another highly inefficient NADPH regeneration cycle, we suggest that MaeB could operate in reverse (pyruvate carboxylating) direction under formatotrophic growth conditions. This carboxylating activity would allow production of biomass precursors (such as oxaloacetate and PEP) through a more efficient route than the alternative via Pps and Ppc. If this is the case, upregulation of MaeB combined with downregulation or knock-out of Pps and Ppc could help promote this more efficient biomass production route.

When comparing the genomes and proteomes of the three formatotrophic strains, we identified some mutations that may contribute to their differences in growth on formate. For instance, we found a mutation in strain K4e in a branching point of the rGlyP with native metabolism (MetF) which may contribute to its improved formatotrophic growth compared to its ancestor K4.

To conclude, our investigation provides insight into the metabolic landscape of engineered and evolved formatotrophic *E. coli* strains, thereby elucidating critical areas for improvement in efficiently utilizing formate as a renewable feedstock for bioproduction (Supplementary Table 5). Our findings offer a roadmap for future rational engineering endeavors aimed at maximizing the efficiency of synthetic formatotrophy. Addressing the identified metabolic bottlenecks and fine-tuning expression targets could pave the way towards industrial applications of formate-based bioproduction systems as sustainable alternative to traditional petrochemical production.

## CRediT authorship contribution statement

**Suzan Yilmaz**: Conceptualization, Data curation, Formal analysis, Investigation, Methodology, Project administration, Visualization, Writing – original draft, Writing – review & editing. **Boas Kanis**: Data curation, Formal analysis, Methodology, Software, Writing – review & editing. **Rensco A. H. Hogers**: Data curation, Formal analysis, Methodology, Software. **Sara Benito-Vaquerizo**: Conceptualization, Data curation, Formal analysis, Methodology, Software, Writing – review & editing. **Jörg Kahnt**: Data curation, Formal analysis, Investigation, Methodology. **Timo Glatter**: Methodology, Writing – review & editing. **Beau Dronsella**: Methodology, Writing – review & editing. **Tobias J. Erb**: Funding acquisition, Supervision. **Maria Suarez-Diez**: Conceptualization, Funding acquisition, Methodology, Project administration, Supervision, Writing – review & editing. **Nico J. Claassens**: Conceptualization, Funding acquisition, Methodology, Project administration, Supervision, Writing – review & editing

## Acknowledgements

We thank Seohyoung Kim, Steffen Lindner, and Arren Bar-Even, for creating and donating the strains used in this study and for their advice and support. We also thank John van der Oost for his advice and support and we thank Vittorio Rainaldi for restoring the AceA gene in the K4 and K4e2 stains that were used in this study. We thank Bart Nijsse for his contributions in data analysis. S.Y. and N.J.C. acknowledge the support of the Dutch Research Council (NWO) via the Gravitation Project BaSyC (024.003.019). In addition, N.J.C. acknowledges support from his NWO Veni fellowship (VI.Veni.192.156).

