## Supplementary Figures and Tables for "System-level characterization of engineered and evolved formatotrophic *E. coli* strains"

### 1.1. Growth data:

Table 1. Overview of doubling times (in hours) of E. coli strains growing via the rGlyP on minimal medium with various concentrations of formate (indicated on the top row in mM). NG: no growth

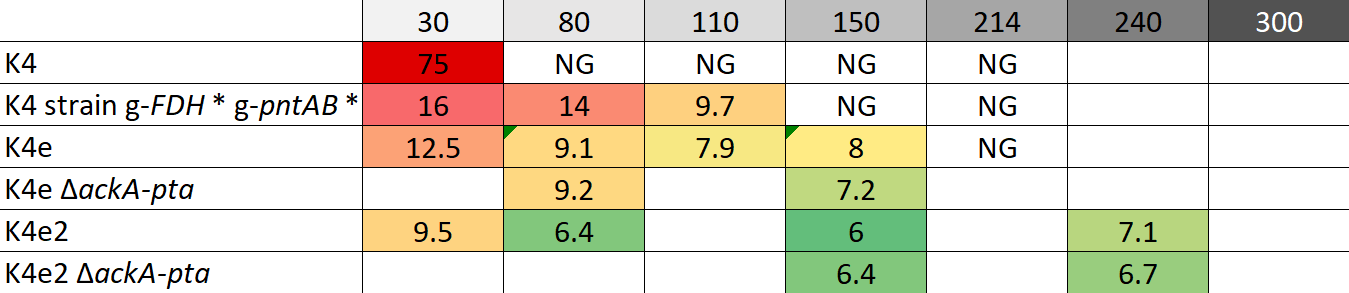

### 1.2. Proteomics data:

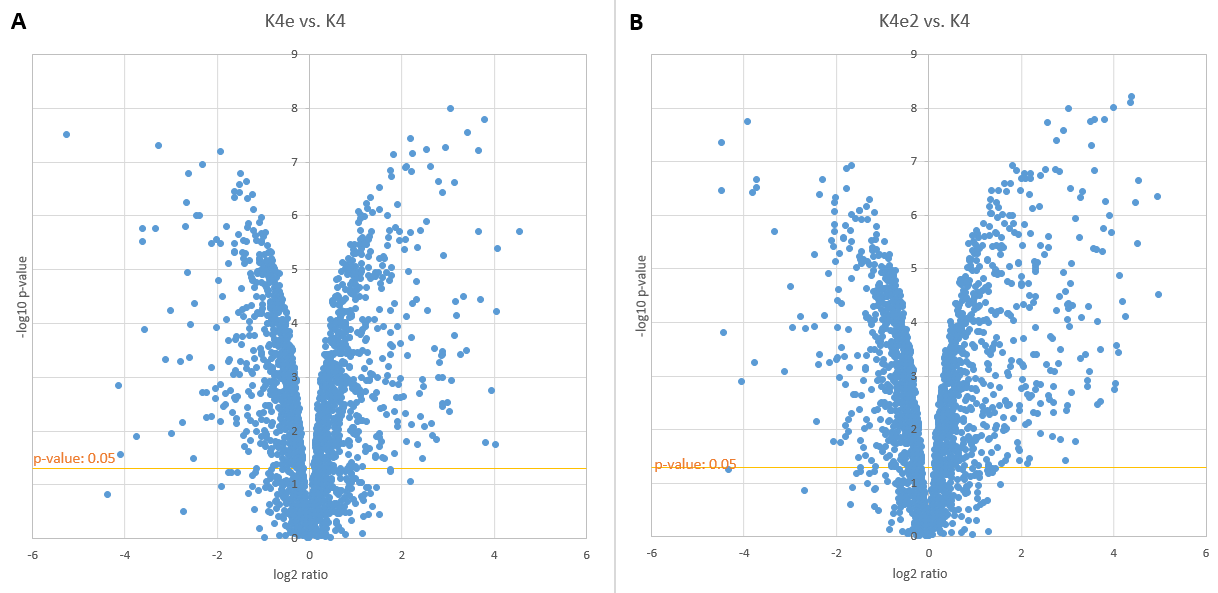

Figure1. Volcano plots of showing abundance differences for all detected proteins. X-axis indicates log2 fold change in protein abundance and y-axis indicates -log10 p-value. (**A**) K4e abundance / K4 abundance. (**B**) K4e2 abundance / K4e abundance

Table 2. Protein groups and protein abundance data used for Figure 9.

|  | **Protein** | **Protein Description** | **K4 formate** | **K4e formate** | **K4e2 formate** | **glucose** |
| --- | --- | --- | --- | --- | --- | --- |
| PsFDH | PsFDH | Formate dehydrogenase | 2.75E-01 | 3.69E+00 | 4.27E+00 | na |
| rGlyP C1 | MeFch | 5,10-methenyl-THF cyclohydrolase | 1.07E-01 | 1.04E-01 | 9.87E-02 | na |
|  | MeFtfL | Formate-THF ligase | 2.18E+00 | 2.03E+00 | 2.35E+00 | na |
|  | MeMtdA | 5,10-methylene-THF dehydrogenase | 1.58E-01 | 1.16E-01 | 1.95E-01 | na |
| rGlyP C2 | GcvH | Glycine cleavage system H protein | 1.35E+00 | 1.51E+00 | 1.85E+00 | 6.01E-01 |
|  | GcvT | Aminomethyltransferase | 1.04E+00 | 1.05E+00 | 1.87E+00 | 1.73E-01 |
|  | GcvP | Glycine dehydrogenase (decarboxylating) | 1.25E+00 | 1.50E+00 | 1.43E+00 | 7.66E-02 |
|  | LpdA | Dihydrolipoyl dehydrogenase | 4.94E-01 | 4.60E-01 | 7.82E-01 | 3.67E-01 |
| rGlyP C3 | SdaA | L-serine dehydratase 1 | 2.04E+00 | 2.21E+00 | 2.16E+00 | 1.15E-01 |
|  | GlyA | Serine hydroxymethyltransferase | 8.70E-01 | 7.77E-01 | 9.45E-01 | 4.53E-01 |
| GNG/  Glycolysis | Glk | Glucokinase | 6.15E-02 | 6.13E-02 | 4.14E-02 | 9.76E-02 |
|  | Pgi | Glucose-6-phosphate isomerase | 5.96E-02 | 4.39E-02 | 4.03E-02 | 5.13E-02 |
|  | PfkA | ATP-dependent 6-phosphofructokinase isozyme 1 | 3.78E-02 | 2.88E-02 | 2.77E-02 | 1.61E-01 |
|  | PfkB | ATP-dependent 6-phosphofructokinase isozyme 2 | 5.51E-02 | 5.13E-02 | 7.06E-02 | 1.78E-01 |
|  | Fbp | Fructose-1,6-bisphosphatase class 1 | 2.80E-02 | 5.21E-02 | 4.23E-02 | 2.56E-02 |
|  | FbaA | Fructose-bisphosphate aldolase class 2 | 2.55E-02 | 1.75E-02 | 2.81E-02 | 8.22E-01 |
|  | FbaB | Fructose-bisphosphate aldolase class 1 | 1.99E-01 | 3.32E-01 | 5.39E-01 | 3.66E-02 |
|  | TpiA | Triosephosphate isomerase | 1.15E-01 | 8.72E-02 | 1.41E-01 | 7.59E-01 |
|  | GlpD | Aerobic glycerol-3-phosphate dehydrogenase | 2.00E-03 | 7.12E-04 | 1.02E-03 | 1.23E-02 |
|  | GapA | Glyceraldehyde-3-phosphate dehydrogenase A | 1.52E-01 | 1.17E-01 | 1.77E-01 | 9.91E-01 |
|  | Epd | D-erythrose-4-phosphate dehydrogenase | 3.85E-03 | 4.15E-03 | 3.44E-03 | 1.42E-01 |
|  | Pgk | Phosphoglycerate kinase | 6.55E-01 | 6.16E-01 | 7.28E-01 | 1.20E+00 |
|  | GpmA | 2,3-bisphosphoglycerate-dependent  phosphoglycerate mutase | 1.68E-01 | 1.31E-01 | 1.62E-01 | 1.03E+00 |
|  | GpmI | 2,3-bisphosphoglycerate-independent  phosphoglycerate mutase | 6.58E-02 | 2.49E-02 | 4.52E-02 | 8.84E-02 |
|  | Eno | Enolase | 2.70E-01 | 2.16E-01 | 2.88E-01 | 1.04E+00 |
| TCA | GltA | Citrate synthase | 3.82E-01 | 6.58E-01 | 3.74E-01 | 1.10E-01 |
|  | GltB | Glutamate synthase [NADPH] large chain | 1.21E-01 | 4.36E-02 | 1.17E-01 | 1.20E-02 |
|  | AcnA | Aconitate hydratase A | 1.38E-01 | 3.46E-01 | 3.37E-01 | 8.67E-02 |
|  | AcnB | Aconitate hydratase B | 1.23E-01 | 5.51E-01 | 4.99E-01 | 4.93E-01 |
|  | Icd | Isocitrate dehydrogenase [NADP] | 2.98E-01 | 3.09E-01 | 3.77E-01 | 8.99E-01 |
|  | SucA | 2-oxoglutarate dehydrogenase E1 component | 8.33E-02 | 1.91E-01 | 1.18E-01 | 6.85E-02 |
|  | SucB | Dihydrolipoyllysine-residue succinyltransferase  component of 2-oxoglutarate dehydrogenase complex | 5.50E-02 | 1.17E-01 | 8.22E-02 | 4.40E-02 |
|  | SucC | Succinate--CoA ligase [ADP-forming] subunit beta | 3.62E-01 | 8.65E-01 | 5.18E-01 | 3.22E-01 |
|  | SucD | Succinate--CoA ligase [ADP-forming] subunit alpha | 3.27E-01 | 7.07E-01 | 4.22E-01 | 8.71E-01 |
|  | SdhA | Succinate dehydrogenase flavoprotein subunit | 2.27E-01 | 5.09E-01 | 2.88E-01 | 1.93E-01 |
|  | SdhB | Succinate dehydrogenase iron-sulfur subunit | 9.49E-02 | 2.09E-01 | 1.05E-01 | 2.96E-01 |
|  | SdhD | Succinate dehydrogenase hydrophobic membrane  anchor subunit | 6.13E-03 | 1.94E-02 | 1.20E-02 | na |
|  | FrdA | Fumarate reductase flavoprotein subunit | 3.01E-02 | 5.17E-02 | 3.63E-02 | 2.18E-02 |
|  | FrdB | Fumarate reductase iron-sulfur subunit | 4.10E-03 | 7.62E-03 | 4.69E-03 | 3.95E-01 |
|  | FrdC | Fumarate reductase subunit C | 2.28E-03 | 3.55E-03 | 1.56E-03 | na |
|  | FumA | Fumarate hydratase class I | 1.00E-01 | 2.08E-01 | 1.43E-01 | 1.34E-01 |
|  | FumB | Fumarate hydratase class I, anaerobic | 2.32E-04 | 7.75E-05 | 8.48E-05 | 1.78E-02 |
|  | FumC | Fumarate hydratase class II | 7.98E-02 | 4.48E-02 | 4.10E-02 | 1.66E-01 |
|  | Mdh | Malate dehydrogenase | 7.32E-01 | 1.22E+00 | 6.39E-01 | 2.16E+00 |
| Glyoxylate  shunt | AceA | Isocitrate lyase | 3.05E+00 | 4.81E+00 | 1.07E-02 | 4.70E+00 |
|  | AceB | Malate synthase A | 2.11E-01 | 6.91E-01 | 1.22E+00 | 9.38E-02 |
|  | GlcB | Malate synthase G | 1.92E-02 | 3.65E-02 | 2.72E-02 | 1.87E-01 |
| PPP | TktA | Transketolase 1 | 1.63E-01 | 2.05E-01 | 2.45E-01 | 6.11E-02 |
|  | TktB | Transketolase 2 | 1.24E-01 | 9.48E-02 | 1.39E-01 | 5.71E-02 |
|  | TalA | Transaldolase A | 2.37E-01 | 2.08E-01 | 1.62E-01 | 1.80E-01 |
|  | TalB | Transaldolase B | 2.64E-01 | 2.04E-01 | 2.38E-01 | 3.24E-01 |
|  | RpiA | Ribose-5-phosphate isomerase A | 3.60E-02 | 3.27E-02 | 3.95E-02 | 5.60E-01 |
|  | RpiB | Ribose-5-phosphate isomerase B | 6.36E-04 | 1.45E-03 | 1.71E-03 | 6.49E-02 |
|  | Rpe | Ribulose-phosphate 3-epimerase | 1.47E-02 | 1.45E-02 | 1.14E-02 | 6.37E-02 |
|  | Zwf | Glucose-6-phosphate 1-dehydrogenase | 8.76E-02 | 4.18E-02 | 3.36E-02 | 1.60E-02 |
|  | Pgl | 6-phosphogluconolactonase | 1.20E-01 | 1.00E-01 | 1.18E-01 | 5.98E-01 |
|  | Gnd | 6-phosphogluconate dehydrogenase, decarboxylating | 1.04E-01 | 8.07E-02 | 1.19E-01 | 1.53E-01 |
|  | AldB | Aldehyde dehydrogenase B | 5.24E-02 | 6.36E-02 | 7.66E-02 | 5.79E-01 |

Table 3. Protein abundance data of respiration complexes.

| **Protein** | **Protein description** | **K4** | **K4e** | **K4e2** | **Glucose** |
| --- | --- | --- | --- | --- | --- |
| Ndh | Type II NADH:quinone oxidoreductase | 2.55E-02 | 1.03E-02 | 3.05E-02 | 2.69E-02 |
| CyoA | Cytochrome bo(3) ubiquinol oxidase subunit 2 | 5.92E-02 | 9.89E-02 | 6.59E-02 | 9.13E-01 |
| CyoB | Cytochrome bo(3) ubiquinol oxidase subunit 1 | 3.60E-02 | 7.13E-02 | 4.44E-02 | na |
| nuoA | NADH-quinone oxidoreductase subunit A | 8.65E-03 | 1.30E-02 | 1.54E-02 | na |
| nuoH | NADH-quinone oxidoreductase subunit H | 2.07E-04 | 4.29E-04 | 2.80E-04 | na |
| nuoK | NADH-quinone oxidoreductase subunit K | 1.01E-03 | 1.43E-03 | 1.12E-03 | na |
| nuoB | NADH-quinone oxidoreductase subunit B | 2.32E-02 | 4.09E-02 | 3.87E-02 | 1.28E-01 |
| nuoC | NADH-quinone oxidoreductase subunit C/D | 5.90E-02 | 9.68E-02 | 7.93E-02 | 4.19E-03 |
| nuoE | NADH-quinone oxidoreductase subunit E | 4.03E-03 | 8.84E-03 | 9.64E-03 | 4.51E-01 |
| nuoF | NADH-quinone oxidoreductase subunit F | 2.38E-02 | 4.51E-02 | 3.74E-02 | 1.77E-02 |
| nuoG | NADH-quinone oxidoreductase subunit G | 5.42E-02 | 1.03E-01 | 9.64E-02 | 5.24E-02 |
| nuoI | NADH-quinone oxidoreductase subunit I | 1.89E-02 | 3.40E-02 | 3.10E-02 | 1.29E-01 |
| cydA | Cytochrome bd-I ubiquinol oxidase subunit 1 | 4.36E-02 | 4.57E-02 | 3.89E-02 | 1.94E-02 |
| cydB | Cytochrome bd-I ubiquinol oxidase subunit 2 | 1.93E-02 | 1.06E-02 | 1.52E-02 | na |
| fdoH | Formate dehydrogenase-O iron-sulfur subunit | 1.25E-02 | 1.15E-02 | 5.31E-03 | 4.11E-03 |
| fdoG | Formate dehydrogenase-O major subunit | 8.06E-03 | 6.00E-03 | 4.37E-03 | 1.69E-01 |
| fdhF | Formate dehydrogenase H | 1.35E-04 | 3.45E-05 | 7.00E-04 | na |
| fdnH | Formate dehydrogenase, nitrate-inducible, iron-sulfur subunit | 3.16E-04 | 1.07E-03 | 6.15E-04 | na |

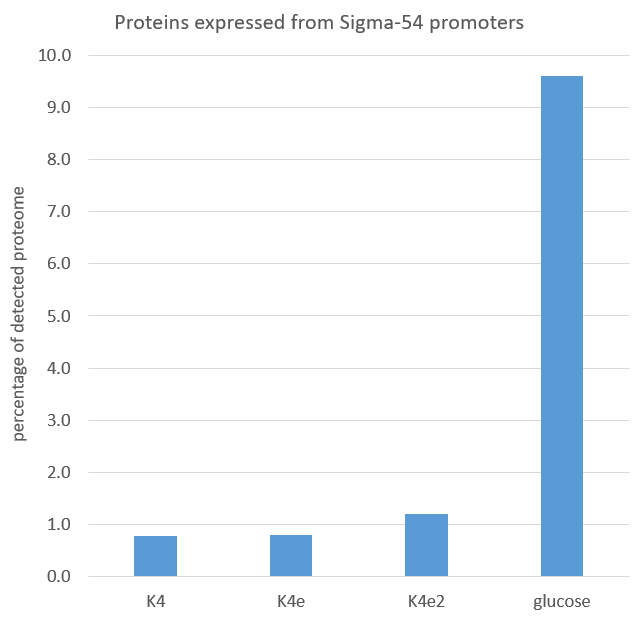

Figure2. Percentages of total quantified proteome for proteins expressed from confirmed [49] sigma-54 promoters. List of proteins and iBAQ data used in Supplementary Table 4.

Table 4. Proteins and iBAQ data used for Supplementary Figure 2.

| **Protein** | **Protein description** | **K4** | **K4e** | **K4e2** | **glucose** |
| --- | --- | --- | --- | --- | --- |
| argT | Lysine/arginine/ornithine-binding  periplasmic protein | 1.1E-01 | 1.6E-01 | 1.6E-01 | 2.6E+00 |
| hisJ | Histidine-binding periplasmic protein | 1.2E-01 | 2.0E-01 | 1.6E-01 | 9.2E-01 |
| hisQ | Histidine transport system permease  protein HisQ | 6.7E-05 | 9.2E-05 | 8.8E-05 | na |
| hisM | Histidine transport system permease  protein HisM | 1.1E-03 | 7.0E-04 | 8.7E-04 | na |
| hisP | Histidine transport ATP-binding protein  HisP | 2.4E-03 | 3.7E-03 | 2.4E-03 | 5.5E-01 |
| astC | Succinylornithine transaminase | 4.7E-04 | 7.8E-04 | 4.7E-04 | 8.2E-01 |
| astA | #N/A | #N/A | #N/A | #N/A | #N/A |
| astD | #N/A | #N/A | #N/A | #N/A | #N/A |
| astB | #N/A | #N/A | #N/A | #N/A | #N/A |
| astE | #N/A | #N/A | #N/A | #N/A | #N/A |
| atoD | #N/A | #N/A | #N/A | #N/A | #N/A |
| atoA | #N/A | #N/A | #N/A | #N/A | #N/A |
| atoE | #N/A | #N/A | #N/A | #N/A | #N/A |
| atoB | #N/A | #N/A | #N/A | #N/A | #N/A |
| fdhF | Formate dehydrogenase H | 1.4E-04 | 3.5E-05 | 7.0E-04 | na |
| glnA | Glutamine synthetase | 3.2E-01 | 2.2E-01 | 4.8E-01 | 3.4E-01 |
| glnL | Sensory histidine kinase/phosphatase NtrB | 2.9E-03 | 1.6E-03 | 2.2E-03 | 2.5E-02 |
| glnG | DNA-binding transcriptional regulator  NtrC | 6.7E-03 | 4.9E-03 | 7.7E-03 | 8.7E-02 |
| glnH | Glutamine-binding periplasmic protein | 4.3E-02 | 8.5E-02 | 7.2E-02 | 1.0E+00 |
| glnP | #N/A | #N/A | #N/A | #N/A | #N/A |
| glnQ | Glutamine transport ATP-binding  protein GlnQ | 1.7E-02 | 2.6E-02 | 2.4E-02 | 4.6E-01 |
| glnK | Nitrogen regulatory protein GlnK | 6.7E-04 | 4.5E-04 | 4.6E-04 | 1.0E-01 |
| amtB | #N/A | #N/A | #N/A | #N/A | #N/A |
| hycA | #N/A | #N/A | #N/A | #N/A | #N/A |
| hycB | Formate hydrogenlyase subunit 2 | 4.2E-04 | 1.2E-04 | 2.6E-03 | na |
| hycC | #N/A | #N/A | #N/A | #N/A | #N/A |
| hycD | #N/A | #N/A | #N/A | #N/A | #N/A |
| hycE | Formate hydrogenlyase subunit 5 | 9.6E-02 | 3.8E-02 | 9.5E-02 | na |
| hycF | Formate hydrogenlyase subunit 6 | 1.0E-04 | 1.5E-04 | 3.8E-04 | na |
| hycG | Formate hydrogenlyase subunit 7 | 2.0E-03 | 1.1E-04 | 8.8E-03 | na |
| hycH | Formate hydrogenlyase maturation  protein HycH | 3.8E-03 | 2.0E-03 | 4.7E-03 | na |
| hycI | Hydrogenase 3 maturation protease | 3.9E-03 | 2.4E-03 | 3.9E-03 | na |
| hydN | Electron transport protein HydN | 3.8E-04 | 1.2E-04 | 5.5E-03 | na |
| hypF | Carbamoyltransferase HypF | 1.6E-03 | 1.0E-03 | 1.0E-03 | na |
| hypA | Hydrogenase maturation factor HypA | 1.4E-03 | 1.6E-04 | 3.3E-03 | na |
| hypB | Hydrogenase maturation factor HypB | 5.2E-03 | 3.0E-03 | 8.9E-03 | 1.2E-02 |
| hypC | Hydrogenase maturation factor HypC | 7.6E-04 | 2.5E-04 | 6.6E-04 | na |
| hypD | Hydrogenase maturation factor HypD | 9.5E-05 | 1.1E-04 | 1.2E-03 | na |
| hypE | Carbamoyl dehydratase HypE | 5.1E-03 | 2.0E-03 | 3.5E-03 | na |
| fhlA | Formate hydrogenlyase transcriptional  activator FhlA | 1.0E-03 | 9.4E-04 | 5.2E-04 | na |
| nac | #N/A | #N/A | #N/A | #N/A | #N/A |
| prpB | 2-methylisocitrate lyase | 1.2E-04 | 3.4E-04 | 2.5E-02 | 4.0E-01 |
| prpC | 2-methylcitrate synthase | 2.0E-04 | 2.5E-04 | 2.0E-02 | 6.8E-02 |
| prpD | 2-methylcitrate dehydratase | 1.6E-03 | 2.9E-03 | 3.5E-02 | 5.8E-02 |
| prpE | Propionate--CoA ligase | 2.2E-04 | 2.6E-04 | 1.1E-03 | na |
| pspA | Phage shock protein A | 2.7E-02 | 2.8E-02 | 3.5E-02 | 1.5E-01 |
| pspB | #N/A | #N/A | #N/A | #N/A | #N/A |
| pspC | Phage shock protein C | 2.0E-03 | 1.9E-03 | 1.4E-03 | na |
| pspD | #N/A | #N/A | #N/A | #N/A | #N/A |
| pspE | Thiosulfate sulfurtransferase PspE | 2.1E-03 | 1.1E-02 | 2.3E-02 | 2.0E+00 |
| rtcB | #N/A | #N/A | #N/A | #N/A | #N/A |
| rtcA | #N/A | #N/A | #N/A | #N/A | #N/A |
| ygiG | #N/A | #N/A | #N/A | #N/A | #N/A |
| zraP | #N/A | #N/A | #N/A | #N/A | #N/A |
| zraS | #N/A | #N/A | #N/A | #N/A | #N/A |
| sraR | #N/A | #N/A | #N/A | #N/A | #N/A |
| hydG | #N/A | #N/A | #N/A | #N/A | #N/A |
| hydH | #N/A | #N/A | #N/A | #N/A | #N/A |

### 1.3. Additional proteins K4, K4e2, and K4e2:

>Ps_Fdh
MHHHHHHAKVLCVLYDDPVDGYPKTYARDDLPKIDHYPGGQTLPTPKAIDFTPGQLLGSVSGELGLRKYLESNGHTLVVTSDKDGPDSVFERELVDADVVISQPFWPAYLTPERIAKAKNLKLALTAGIGSDHVDLQSAIDRNVTVAEVTYCNSISVAEHVVMMILSLVRNYLPSHEWARKGGWNIADCVSHAYDLEAMHVGTVAAGRIGLAVLRRLAPFDVHLHYTDRHRLPESVEKELNLTWHATREDMYPVCDVVTLNCPLHPETEHMINDETLKLFKRGAYIVNTARGKLCDRDAVARALESGRLAGYAGDVWFPQPAPKDHPWRTMPYNGMTPHISGTTLTAQARYAAGTREILECFFEGRPIRDEYLIVQGGALAGTGAHSYSKGNATGGSEEAAKFKKAV

>Me_FtfL
MHHHHHHPSDIEIARAATLKPIAQVAEKLGIPDEALHNYGKHIAKIDHDFIASLEGKPEGKLVLVTAISPTPAGEGKTTTTVGLGDALNRIGKRAVMCLREPSLGPCFGMKGGAAGGGKAQVVPMEQINLHFTGDFHAITSAHSLAAALIDNHIYWANELNIDVRRIHWRRVVDMNDRALRAINQSLGGVANGFPREDGFDITVASEVMAVFCLAKNLADLEERLGRIVIAETRDRKPVTLADVKATGAMTVLLKDALQPNLVQTLEGNPALIHGGPFANIAHGCNSVIATRTGLRLADYTVTEAGFGADLGAEKFIDIKCRQTGLKPSAVVIVATIRALKMHGGVNKKDLQAENLDALEKGFANLERHVNNVRSFGLPVVVGVNHFFQDTDAEHARLKELCRDRLQVEAITCKHWAEGGAGAEALAQAVVKLAEGEQKPLTFAYETETKITDKIKAIATKLYGAADIQIESKAATKLAGFEKDGYGGLPVCMAKTQYSFSTDPTLMGAPSGHLVSVRDVRLSAGAGFVVVICGEIMTMPGLPKVPAADTIRLDANGQIDGLF

>Me_Fch
MHHHHHHAGNETIETFLDGLASSAPTPGGGGAAAISGAMGAALVSMVCNLTIGKKKYVEVEADLKQVLEKSEGLRRTLTGMIADDVEAFDAVMGAYGLPKNTDEEKAARAAKIQEALKTATDVPLACCRVCREVIDLAEIVAEKGNLNVISDAGVAVLSAYAGLRSAALNVYVNAKGLDDRAFAEERLKELEGLLAEAGALNERIYETVKSKVN

>Me_MtdA
MHHHHHHSKKLLFQFDTDATPSVFDVVVGYDGGADHITGYGNVTPDNVGAYVDGTIYTRGGKEKQSTAIFVGGGDMAAGERVFEAVKKRFFGPFRVSCMLDSNGSNTTAAAGVALVVKAAGGSVKGKKAVVLAGTGPVGMRSAALLAGEGAEVVLCGRKLDKAQAAADSVNKRFKVNVTAAETADDASRAEAVKGAHFVFTAGAIGLELLPQAAWQNESSIEIVADYNAQPPLGIGGIDATDKGKEYGGKRAFGALGIGGLKLKLHRACIAKLFESSEGVFDAEEIYKLAKEMA

### 1.4. Proteomaps:

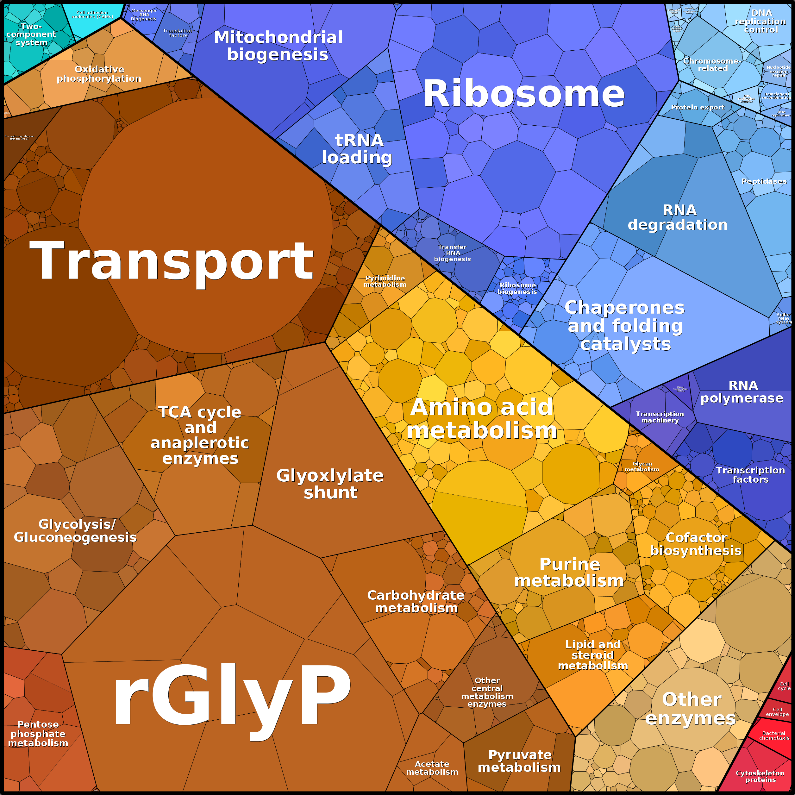

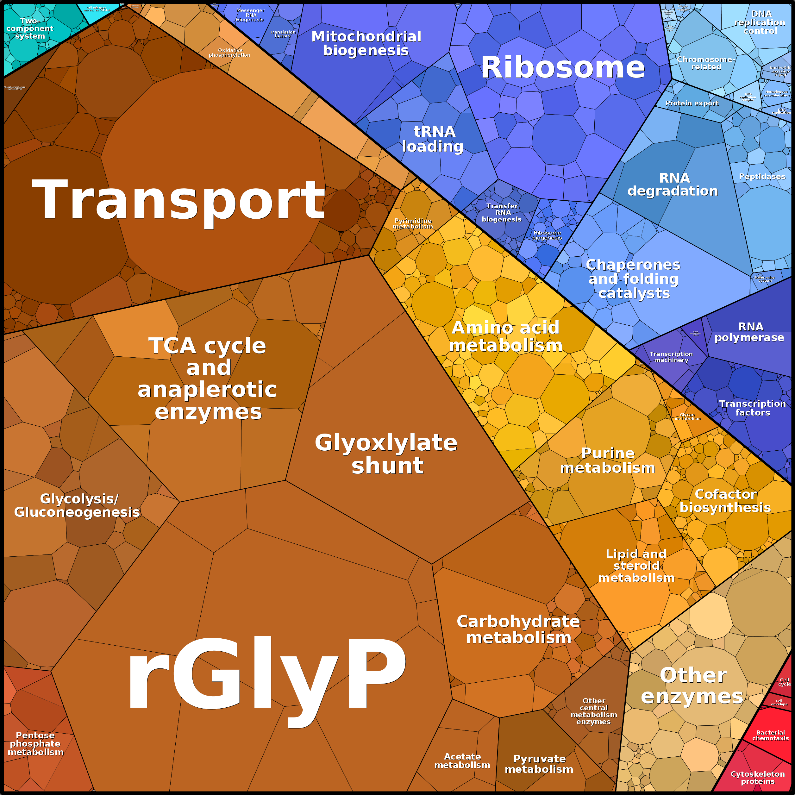

 **K4	K4e**

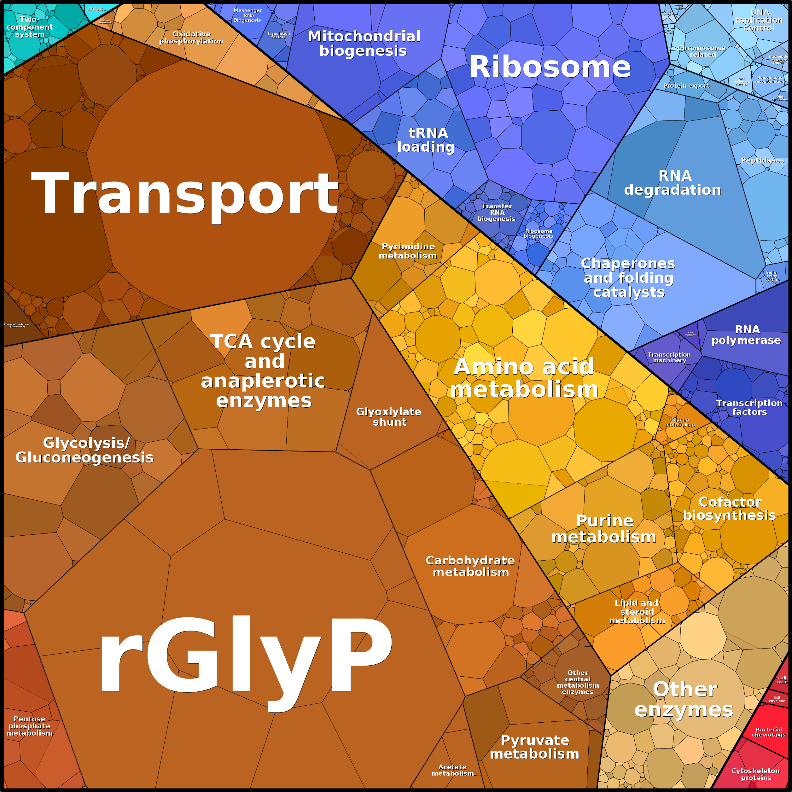

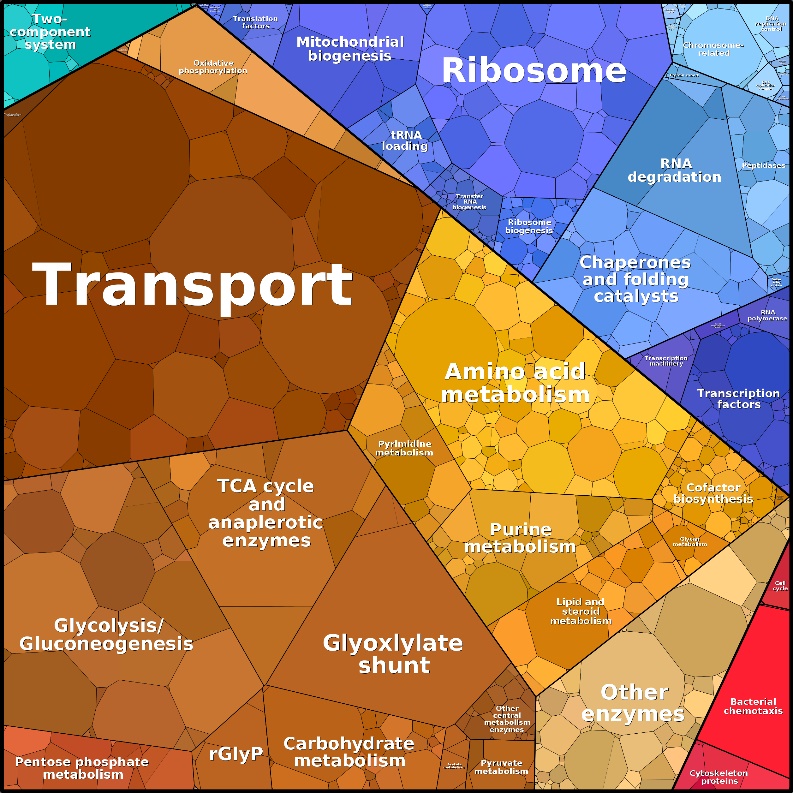

 **K4e2	WT glucose**

Figure 3. Proteomaps displaying categories of proteins based on cellular function proportional to the size of their proteome fraction. Proteomaps were generated using the tool developed by [58], [59], [60].

### 1.5. Suggested mutation targets:

Table 5. Overview of mutations suggested in this article.

| **Pathways** | **Targets** | **Types of mutations** | **Alternative suggestions** |
| --- | --- | --- | --- |
| NADH regeneration | PsFDH | Upregulation | Knock in of faster FDH  such as CnFDH |
| NADPH regeneration | PntAB | Upregulation | Knock in of NADPH  regenerating FDH |
|  | AdhE AldB Acs | Knock out |  |
|  | Zwf | Knock out or knock down |  |
| ATP regeneration | NuoABCEFGHIJKLMN,  CyoABCD | Expression tuning? |  |
|  | other respiratory complexes  (Ndh, CydABHX, AppBCX,  FdnGHI, FdoGHI) | Knock down? |  |
| Oxidative TCA cycle | SucAB | Knock out or knock down |  |
|  | GltA | Knock down |  |
| Oxaloacetate/PEP production | MaeB, PpsA, Ppc | Expression tuning |  |
| Glyoxylate shunt | (AceA), AceB | Knock out or knock down |  |
| Acetate production | PoxB | Knock out |  |
| PDH and GCV | AceE, AceF | Knock down |  |
|  | Lpd | Upregulation |  |
